# Sex differences in IL-17 determine chronicity in male versus female urinary tract infection

**DOI:** 10.1101/449124

**Authors:** Anna Zychlinsky Scharff, Matthieu Rousseau, Livia Lacerda Mariano, Tracy Canton, Matthew L Albert, Magnus Fontes, Darragh Duffy, Molly A Ingersoll

## Abstract

Sex-based differences influence incidence and outcome of infectious disease. Women have a significantly greater incidence of urinary tract infection (UTI) than men, yet, conversely, male UTI is more persistent with greater associated morbidity. Mechanisms underlying these sex-based differences are unknown, in part due to a lack of experimental models. We optimized a model to transurethrally infect male mice and directly compared UTI in both sexes. Although both sexes were initially equally colonized by uropathogenic *E. coli*, only male and testosterone-treated female mice remained chronically infected for up to 4 weeks. Female mice had more robust innate responses, including higher IL-17 expression, and increased γδ T and LTi-like cells in the bladder following infection. Accordingly, neutralizing IL-17 abolished resolution in female mice, identifying the cytokine pathway necessary for bacterial clearance. Our findings support the concept that sex-based responses to UTI contribute to impaired innate immunity in males and provide a rationale for non-antibiotic-based immune targeting to improve the response to UTI.

**One Sentence Summary:** We investigated mechanisms underlying the clinical observation that while urinary tract infection is more prevalent in women, it is more severe in men, observing that in contrast to robust immune responses characterized by IL-17 in female animals, male mice develop chronic infection, following a failure to initiate innate immunity.

## Introduction

Mounting evidence points to the importance of sex in determining susceptibility to infection and immune responses (Ingersoll, 2017; Klein and Flanagan, 2016; vom Steeg and Klein, 2016). For instance, sex-specific differences are evident in bacterial infections by *Legionella, Mycobacterium*, and *Campylobacter* (vom Steeg and Klein, 2016). Urinary tract infections (UTI) exhibit one of the most distinctive sex differences among infectious diseases; adult premenopausal women are 40 times more likely than adult men to experience UTI (Foxman, 2010). Consequently, basic and clinical research has overwhelmingly focused on the pathogenesis of UTI in females with little regard for the underlying reasons for this disparity. Importantly, this bias in adult women is not representative of the overall prevalence of UTI in the general population. Indeed, the incidence of UTI is similar between the sexes in infants and the elderly (Harper and Fowlis, 2007). For example, 40% of pediatric UTI patients (<24 months of age) are male (Craig et al., 2009). Similarly, the incidence of UTI was 14% in women and 11% in men in a healthy geriatric population over the age of 65 (Ruben et al., 1995). These changes in incidence suggest that factors such as sex hormone levels might influence susceptibility to infection during a lifetime (Ingersoll, 2017). Notably, although the incidence of male UTI in adulthood is significantly lower compared to adult women, male UTI patients have an elevated risk of morbidity from complications, as infection in men of all ages is clinically classified as a complicated UTI (Conway et al., 2015; Fabbian et al., 2015; Lipsky, 1989). Indeed, although the same first-line antibiotics are used in female and male patients, men require longer treatment durations to eradicate bacteria (Tandan et al., 2016). Together, increased severity and treatment challenges, combined with mounting antibiotic resistance in uropathogenic *E. coli* (UPEC) strains, highlight the need to better understand immunity to UTI in women and men, to determine the mechanisms mediating disease outcome in both sexes. In turn, this knowledge may contribute to the development of non-antibiotic based stratified therapies targeting sex-specific immune pathways for the treatment of women and men with UTI.

Given the frequency and high morbidity of UTI in infant and elderly males, it is disconcerting that our knowledge of UTI pathogenesis is based almost entirely on studies in female animals. Indeed, transurethral instillation models used to study UTI are performed almost exclusively in female rodents (Balemans et al., 1994; Bjorling et al., 2011; Hung et al., 2009), and the technique is broadly reported to be unachievable in male animals (El Behi et al., 2013; Hagberg et al., 1983; Oliveira et al., 2009; Olson et al., 2015; Seager et al., 2009). Only a small number of laboratories report the use of transurethral instillation of bacteria into the male urogenital tract. Studies from these groups investigate prostatitis, and no report has included quantification of bacterial colonization in the bladder or comparison to infected female animals (Boehm et al., 2012; Elkahwaji et al., 2007; Lee et al., 2015; Shinohara et al., 2013; Simons et al., 2015; Wong et al., 2014, 2015). To date, only one animal study explored sex differences in UTI, by surgically exposing the bladder abdominally and injecting bacteria through the bladder wall to establish infection (Olson et al., 2016). The authors observed that male C3H/HeN mice, but not C57BL/6J mice, remain chronically infected following infection (Olson et al., 2016). Importantly, however, in this model it cannot be excluded that surgery-related inflammation and secondary effects of wound-healing pathways, known to differ between the sexes, influenced bacterial clearance (Emmerson et al., 2009; Gilliver et al., 2007; Knipper et al., 2015).

Thus, to test the hypothesis that divergent host responses to UTI between male and female animals results in differential outcomes following infection with UPEC, we established a method of intravesical instillation of bacteria *via* catheterization of the urethra of male mice (Zychlinsky Scharff et al., 2017). This protocol, derived from one routinely used for female UTI studies (Hung et al., 2009), permitted direct comparison of the innate and adaptive immune response in infected female and male animals over time. C57BL/6J female mice resolved their infection without intervention, as previously reported (Mora-Bau et al., 2015; Mysorekar and Hultgren, 2006), however male mice displayed persistent bacteriuria, remaining chronically infected for at least one month. Within the first 24 hours following infection, female mice exhibited more robust cytokine expression accompanied by greater immune cell infiltration. Cytokine expression, immune cell infiltration, and resolution were attenuated or abrogated in testosterone-treated females. These findings suggest that the increased severity associated with male UTI is due to hormone-mediated suppression of the innate immune response.

## Results

### Male mice fail to resolve UPEC infection

To directly compare transurethrally infected female and male animals, we adapted a protocol to catheterize female mice to male animals (Hung et al., 2009). A detailed video protocol of our method is reported elsewhere (Zychlinsky Scharff et al., 2017). To investigate the capacity of female and male mice to resolve UPEC infection, we intravesically instilled 7 week old C57BL/6J mice of both sexes with 10^7^ colony forming units (CFU) of one of two isogenic UPEC UTI89 strains bearing resistance to ampicillin (UTI89-GFP-amp^R^) or kanamycin (UTI89-RFP-kan^R^) (Mora-Bau et al., 2015). Strikingly, we observed that a majority of male mice remained chronically infected up to one month, as determined by the presence of UPEC in the urine, in contrast to female mice, which resolved infection (**Fig. 1A**). We sacrificed experimental groups at predetermined timepoints to quantify bladder bacterial burden. While female and male mice were equally colonized at 24 and 48 hours post-infection (PI), bacterial burden decreased over time in female mice but remained elevated in male mice, resulting in statistically significantly higher bacterial burdens in male mice 28 days PI (**Fig. 1B**). At 28 days PI, 100% of female mice (21/21), but only 29% (7/24) of male mice had sterile urine (percentages calculated from animals shown in **Fig. 1B**). Importantly, female mice that resolve an acute infection have no UPEC in their urine but maintain bacterial burdens of approximately 10^3^ CFU in quiescent tissue reservoirs in C57BL/6J mice (**Fig. 1B**, 28 days PI) (Mora-Bau et al., 2015; Mysorekar and Hultgren, 2006). In a female UTI model, infection is not confined to the bladder and the kidneys can be colonized by vesicoureteral reflux (Hopkins et al., 1995). To determine the extent of infection in male mice, we infected mice with 10^7^ CFU UTI89-GFP-kan^R^ and assessed bacterial colonization 60-90 minutes PI. Bacterial CFU in male mice were comparable to CFU observed in female mice at the same timepoint (Ingersoll et al., 2008). UPEC also colonized the prostate, kidneys, seminal vesicles, preputial glands, and testes in a majority of mice at this timepoint (**Fig. S1A**). Over time, kidney colonization was equivalent between the sexes until 14 days PI, when CFU declined in female mice and increased in male mice (**Fig. S1B**). Finally, in male mice, the prostate, seminal vesicles, preputial glands, and testes were colonized for at least 14 days (**Fig. S1C**).

**Figure 1.**
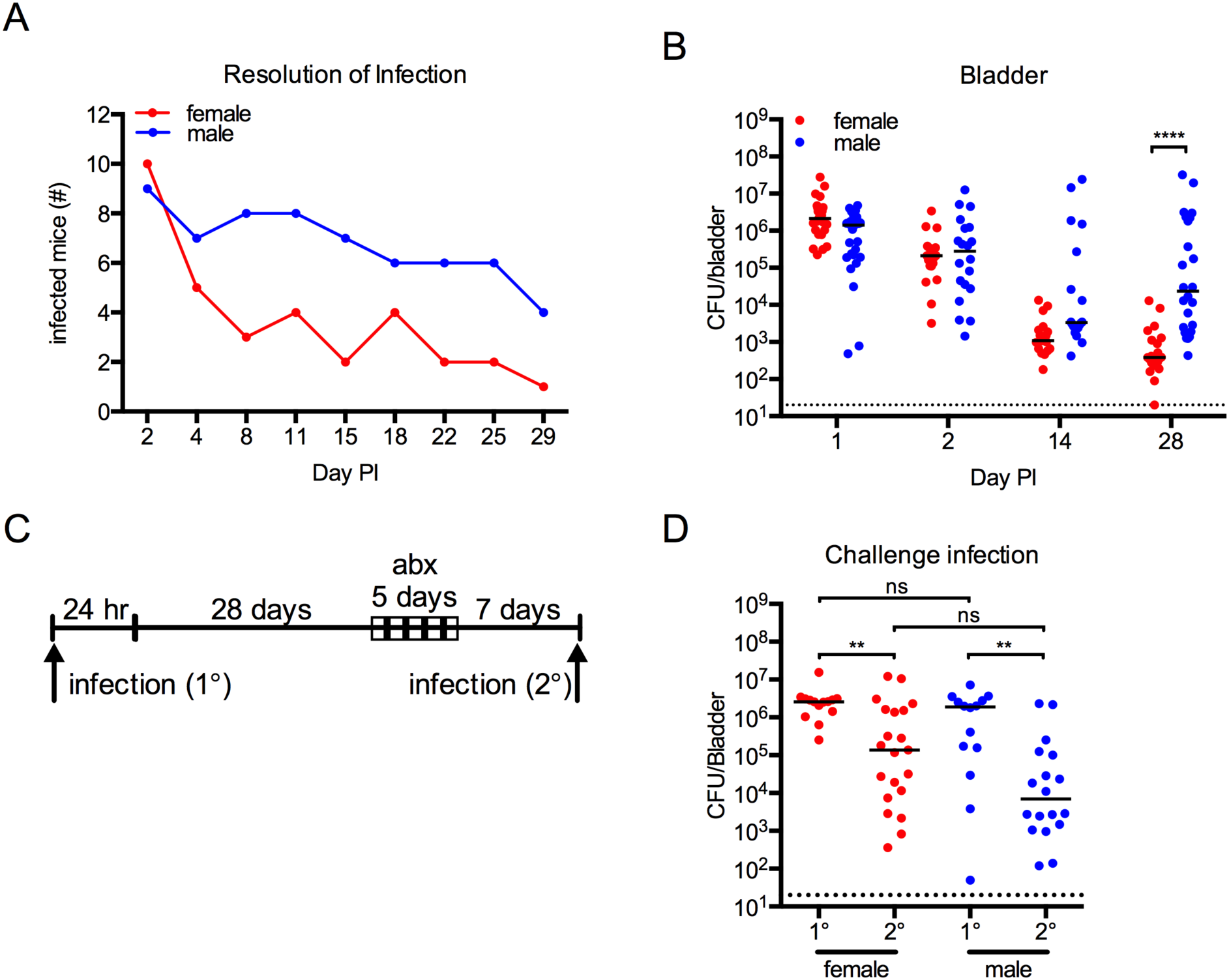
Male mice do not resolve UTI following intravesical instillation. **A-B** 7 week old female and male C57BL/6J mice were infected with 10^7^ CFU of UPEC strain UTI89-RFP-kan^R^ or UTI89-GFP-amp^R^. Graphs depict (**A**) the number of infected mice, determined by urine sampling, and (**B**) CFU/bladder. (**C**) Schematic of the experiment depicted in **D**: Female and male C57BL/6J mice were infected with 10^7^ CFU of UPEC strain UTI89-GFP-amp^R^ and bladder bacterial burden was determined 24 hours post primary (1°) infection. At 29 days post-infection (PI), all animals were treated with antibiotics for 5 days, followed by a 7 day wash-out period. After determination that all mice had sterile urine, mice were challenged with 10^7^ CFU UPEC strain UTI89-RFP-kan^R^ and CFU in bladders determined at 24 hours post challenge (2°) infection. (**D**) The graph shows bacterial CFU at 24 hours post primary or challenge infection in the bladder. **A** depicts representative data from 1 (n=10 female, n=9 male) of more than 5 experiments, red is female, blue is male. Data in **B** and **D**are pooled from 2-4 experiments, n=6-10 mice/group in each experiment. Each dot represents one mouse, red denotes female mice and blue indicates male mice, lines are medians. Dotted line depicts the limit of detection of the assay, 20 CFU/bladder. In **B**, **** p<0.0001, Kruskal-Wallis test comparing female to male at each timepoint, with Dunn’s post-test to correct for multiple comparisons. In **D**, ** p<0.01, ns = not significant, Kruskal-Wallis test comparing 1° to 2° infection within a single sex and comparing 1° or 2° CFU between sexes, with Dunn’s post-test to correct for multiple comparisons.

### Female and male mice mount a nonsterilizing adaptive immune response to UTI

We previously reported that female mice develop a nonsterilizing adaptive immune response following primary UTI that is abrogated in female *Rag2*^−/−^ mice (Mora-Bau et al., 2015). Whereas bacterial burden is reduced 2-3 logs in wildtype mice following challenge infection compared to primary UTI, bacterial burden in *Rag2*^−/−^ mice is equivalent following primary and challenge infection. Indeed, female *Rag2*^−/−^ animals can resolve a primary UTI, suggesting that adaptive immunity is dispensable for bacterial clearance (Mora-Bau et al., 2015). Based on this observation, it was unlikely that a failure to mount an adaptive immune response could explain why chronic infection developed in male mice. To experimentally rule out this possibility, we employed a model of challenge infection (Mora-Bau et al., 2015). Female and male mice were infected with 10^7^ CFU of UTI89-GFP-amp^R^ and a cohort of each sex was sacrificed at 24 hours PI. Although 20/21 female animals had sterile urine, compared to only 3/18 male mice, we treated both sexes beginning at 29 days PI with trimethoprime-sulfamethoxazole (TMP-SMX, Avemix) in the drinking water, to control for potential influence of antibiotic treatment (**Fig. 1C**). Following a 5 day antibiotic treatment and a subsequent 7 day wash-out period, mice were confirmed to have sterile urine and then infected with the isogenic UPEC strain, UTI89-RFP-kan^R^. Comparable to that observed in **Fig. 1B**, bacterial burden was not different between female and male mice 24 hours post primary infection (**Fig. 1D**). As we previously reported, bacterial burden was significantly lower in female mice 24 hours post challenge infection compared to primary infection, and as hypothesized, bacterial burden was also significantly reduced in male mice following challenge infection (**Fig. 1D**). Bacterial burden in challenged animals was not statistically significantly different between the sexes, suggesting that both female and male mice mount an adaptive immune response capable of reducing but not eliminating bacterial burden following a challenge infection.

### Female mice exhibit greater cytokine responses

The results of our challenge infection experiments ruled out the possibility that a weak or absent adaptive immune response accounted for differences in resolution of UTI between the sexes. In addition, as female *Rag2*^−/−^ mice, lacking T and B cells, or T cell depleted mice resolve primary infection with the same kinetic as wildtype female mice (Mora-Bau et al., 2015), we reasoned that the failure to resolve infection in male mice was due to differences in an innate mechanism. To identify innate host factors contributing to resolution in female mice or chronic infection in male mice, we assessed cytokine expression at 24 and 48 hours PI, which is the peak of the cytokine response in female mice (Ingersoll et al., 2008). Female and male mice were instilled with PBS or infected with UPEC strain UTI89-GFP-amp^R^ or UTI89-RFP-kan^R^ as above, and 32 cytokines were analyzed using Luminex multianalyte analysis on whole bladder tissue homogenates at 24 and 48 hours PI. Twenty-six cytokines were expressed above the minimum detectable concentration plus 2 standard deviations in bladders in at least one condition. 23/26 analytes measured were significantly different (T-test, false discovery rate adjusted p value: q<0.05) between PBS-instilled and UPEC-infected female mice at 24 hours PI (**Table 1**). By contrast, only 7/26 analytes were statistically significantly different between control PBS-treated and UPEC-infected male animals (**Table 1**, T-test, q<0.05). ANOVA revealed that 21/26 analytes were significantly differentially expressed among the four groups of PBS-treated and UPEC-infected female and male mice (**Table 1**, q<0.05). Unsupervised hierarchical clustering of the cytokine expression data revealed that UPEC-infected females exhibited a robust inflammatory signature 24 hours PI, whereas infected male mice clustered more closely to PBS-treated animals of both sexes than infected female mice (**Fig. 2A**). Absolute protein expression levels of the 23 cytokines from **Table 1** (PBS *vs.* UPEC-infected female mice) were, overall, greater in infected female animals compared to infected male mice and PBS-treated mice of both sexes at 24 hours PI (**Fig. 2B**). This differential response was short-lived, as female and male mice had, with a few notable exceptions, similar cytokine levels at 48 hours PI, which approached levels comparable to those measured in naïve, untreated bladders (**Fig. 2C**). These differences in cytokine expression were unexpected, as bacterial colonization was equivalent at 24 and 48 hours PI between the sexes (**Fig. 1B**).

**Figure 2.**
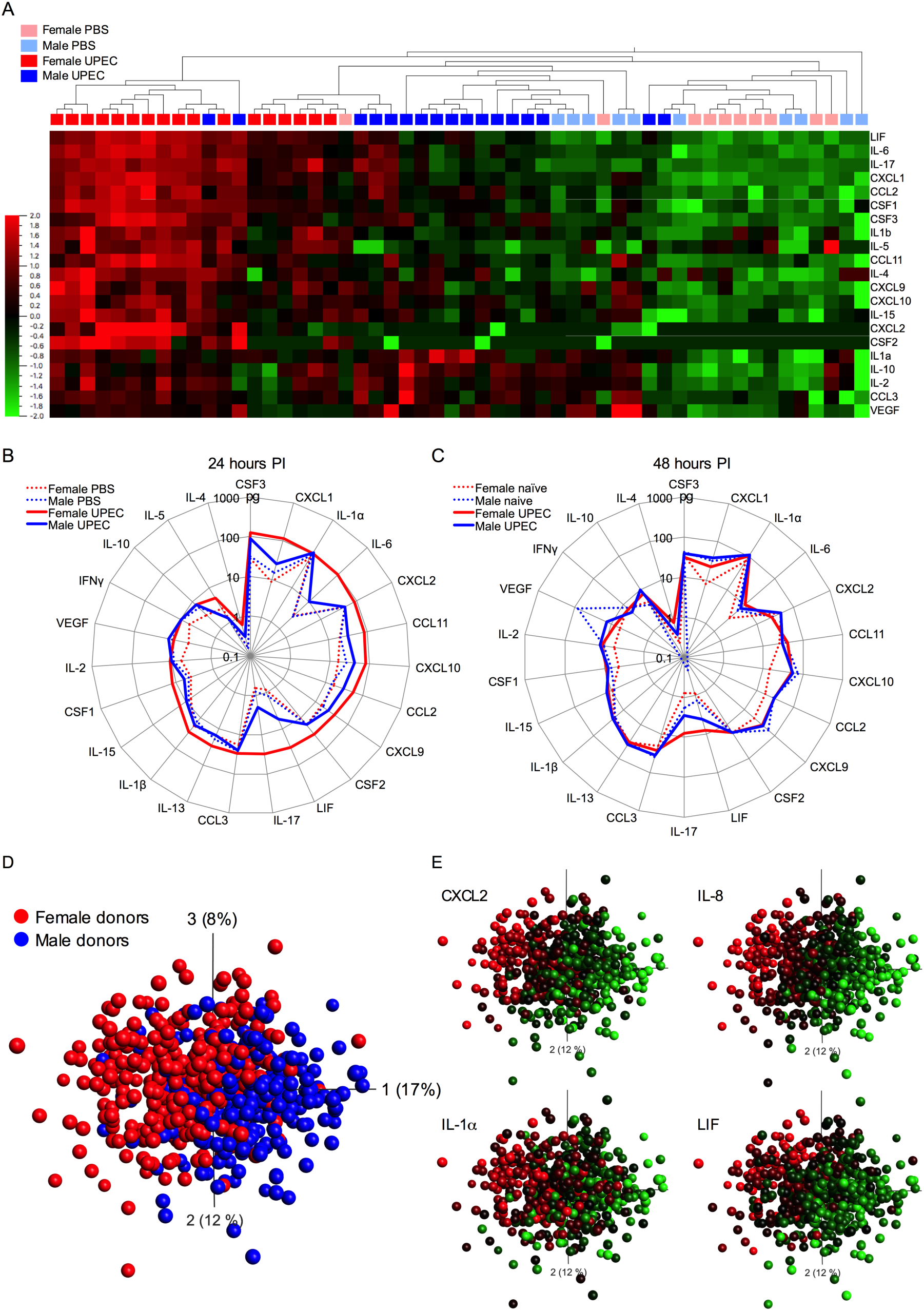
Female mice exhibit a more rapid and robust cytokine response to UTI. Female and male mice were instilled with PBS or infected with 10^7^ CFU of UPEC strain UTI89-RFP-kan^R^ or UTI89-GFP-amp^R^ and sacrificed 24 or 48 hours PI. 32 cytokines were measured in homogenized bladders by Luminex assay. (**A**) Heat map shows unsupervised hierarchical clustering of the 21 significantly different analytes determined by ANOVA (false discovery rate (FDR) adjusted p value to correct for multiple testing; q<0.05) among PBS-treated and UPEC-infected female and male mice. (**B**-**C**) Spider plots show absolute cytokine expression levels (pg/mL) on a log-scale at (**B**) 24 hours PI in PBS-treated (dotted lines) and UPEC-infected female (solid red lines) and male (solid blue lines) mice and (**C**) 48 hours PI (solid lines) or naïve, untreated animals (dotted lines). Data are pooled from 4 experiments, n=5-7 mice per experiment. Analytes and p and q values are listed in **Table 1.** (**D-E**) Principal component analysis (PCA) was performed on a data set of the transcriptional response following whole blood stimulation with heat-killed *E. coli* in 600 healthy donors. (**D**) The PCA plot represents the 275 genes that are differently expressed between women (red dots, each dot is a donor) and men (blue dots, each dot is a donor) as determined by ANOVA, q<0.01. (**E**) mRNA expression level heat map PCA overlay for CXCL2, IL-8, IL-1α, and LIF.

**Table 1:** Analytes defining the response to infection, compared to PBS instillation, at 24 hours.

| Analyte | PBS vs UTI Female<br>mice<sup>*#</sup> |  | PBS vs UTI Male<br>mice<sup>‡</sup> |  | PBS vs UTI Both sexes<sup>°</sup> |  |
| --- | --- | --- | --- | --- | --- | --- |
|  | p-value | q-value | p-value | q-value | p-value | q-value |
| LIF | 3.17E-12 | 8.25E-11 | 2.51E-05 | 6.52E-04 | 6.82E-17 | 1.77E-15 |
| CXCL1 | 7.26E-11 | 9.44E-10 | 1.25E-02 | 4.64E-02 | 4.63E-12 | 3.01E-11 |
| IL-17 | 1.13E-10 | 9.79E-10 | 2.81E-03 | 1.46E-02 | 1.80E-12 | 1.56E-11 |
| IL-6 | 7.79E-10 | 5.06E-09 | 1.38E-04 | 1.79E-03 | 7.68E-14 | 9.98E-13 |
| CSF1 | 3.13E-09 | 1.63E-08 |  |  | 7.94E-11 | 3.44E-10 |
| CCL2 | 3.97E-08 | 1.72E-07 | 2.19E-04 | 1.90E-03 | 3.16E-11 | 1.64E-10 |
| CCL11 | 5.06E-07 | 1.88E-06 |  |  | 6.65E-07 | 1.73E-06 |
| CSF3 | 1.07E-06 | 3.49E-06 | 6.35E-04 | 4.13E-03 | 1.73E-08 | 6.41E-08 |
| CXCL10 | 2.56E-06 | 7.38E-06 |  |  | 7.76E-05 | 1.44E-04 |
| VEGF | 6.51E-06 | 1.69E-05 |  |  | 1.44E-02 | 1.79E-02 |
| IL1α | 2.51E-05 | 5.93E-05 | | | 3.18E-04 | 4.86E-04 |
| CXCL9 | 3.88E-05 | 8.41E-05 |  |  | 6.73E-05 | 1.44E-04 |
| IL-15 | 6.04E-05 | 1.21E-04 |  |  | 7.51E-05 | 1.44E-04 |
| IL-4 | 1.02E-04 | 1.90E-04 |  |  | 1.43E-05 | 3.39E-05 |
| IL-2 | 2.24E-04 | 3.89E-04 |  |  | 2.18E-03 | 3.02E-03 |
| IL-10 | 6.57E-04 | 1.07E-03 |  |  | 2.21E-03 | 3.02E-03 |
| CCL3 | 9.38E-04 | 1.36E-03 |  |  | 1.25E-02 | 1.63E-02 |
| IL1β | 8.93E-04 | 1.36E-03 | 9.90E-03 | 4.29E-02 | 1.10E-07 | 3.58E-07 |
| CSF2 | 2.05E-03 | 2.80E-03 |  |  | 1.29E-04 | 2.10E-04 |
| CXCL2 | 3.34E-03 | 4.34E-03 |  |  | 8.86E-05 | 1.54E-04 |
| IL-13 | 1.58E-02 | 1.96E-02 |  |  |  |  |
| IL-5 | 2.30E-02 | 2.72E-02 |  |  | 1.87E-07 | 5.39E-07 |
| IFNγ | 3.20E-02 | 3.62E-02 | | | 4.66E-04 | 8.65E-04 |

\*Values are ordered by decreasing q-value for analytes that are statistically significantly different between PBS- and UPEC-instilled female mice. q-values are the false discovery rate adjusted p value.

To determine whether this distinct pattern of cytokine expression might be relevant in the context of human infection, we took advantage of a dataset from a recently published study describing the impact of sex on transcriptional variation in induced human immune responses (Piasecka et al., 2018). We investigated differences in gene expression between woman and men aged 20-49 (to avoid potential cofounding effects in post-menopausal women), following whole blood stimulation with *E. coli*. We observed that 49% (275/560) of the genes measured showed a significant (n=600, q<0.01) sex association (**Fig. 2D, Table S1**). The expression of CXCL2, IL-8, IL-1α, and LIF (**Fig. 2E**), as well as CCL3, IL-4, and IL1β (Table S1), were all significantly (q<0.01) higher in female donors after *E. coli* stimulation. A sex-association with the expression of IL-4R and IL-13R, as well as other receptors for cytokines differentially expressed in our mouse model, was also observed (**Table S1**), which would be expected to influence how circulating cells expressing these receptors respond to their cognate cytokines. Together, these findings directly support the potential clinical relevance of our findings in this mouse model.

### Immune cell infiltration is greater in infected female mice

Many of the cytokines induced during UTI facilitate the recruitment of innate immune cells to the site of infection (Hang et al., 1999). To test whether the divergent cytokine response between the sexes impacted immune cell infiltration, we infected female and male mice with UPEC strain UTI89-RFP-kan^R^ and analyzed bladder tissue by flow cytometry at 24 and 48 hours PI. Infected bladders from female mice had significantly greater numbers of total CD45+ immune cells compared to infected male mice (**Fig. 3A**), correlating with increased cytokine expression at this timepoint (**Fig. 2A-B**). Similar to cytokine expression, this difference was evident only at 24 hours PI. While tissue-resident macrophages were not different between the sexes or between naïve and infected animals at these early timepoints, female mice had significantly higher numbers of dendritic cells (DCs) at 24 hours PI (**Fig. 3B**). This increase was modest and did not account for the overall increase in CD45+ immune cell populations. By far the most numerous immune cells were infiltrating neutrophils, monocyte-derived cells, and eosinophils, which were all significantly increased in female mice compared to male mice, only at 24 hours PI (**Fig. 3C, Fig. S2A-B**, gating strategy). Differences in immune cell numbers were not due to baseline differences in naïve bladder resident immune cell populations, as flow cytometric analysis of uninfected bladder tissue revealed that the immune cell compartment was not different between female and male mice (**Fig. 3A-C**, 0 hours PI).

**Figure 3.**
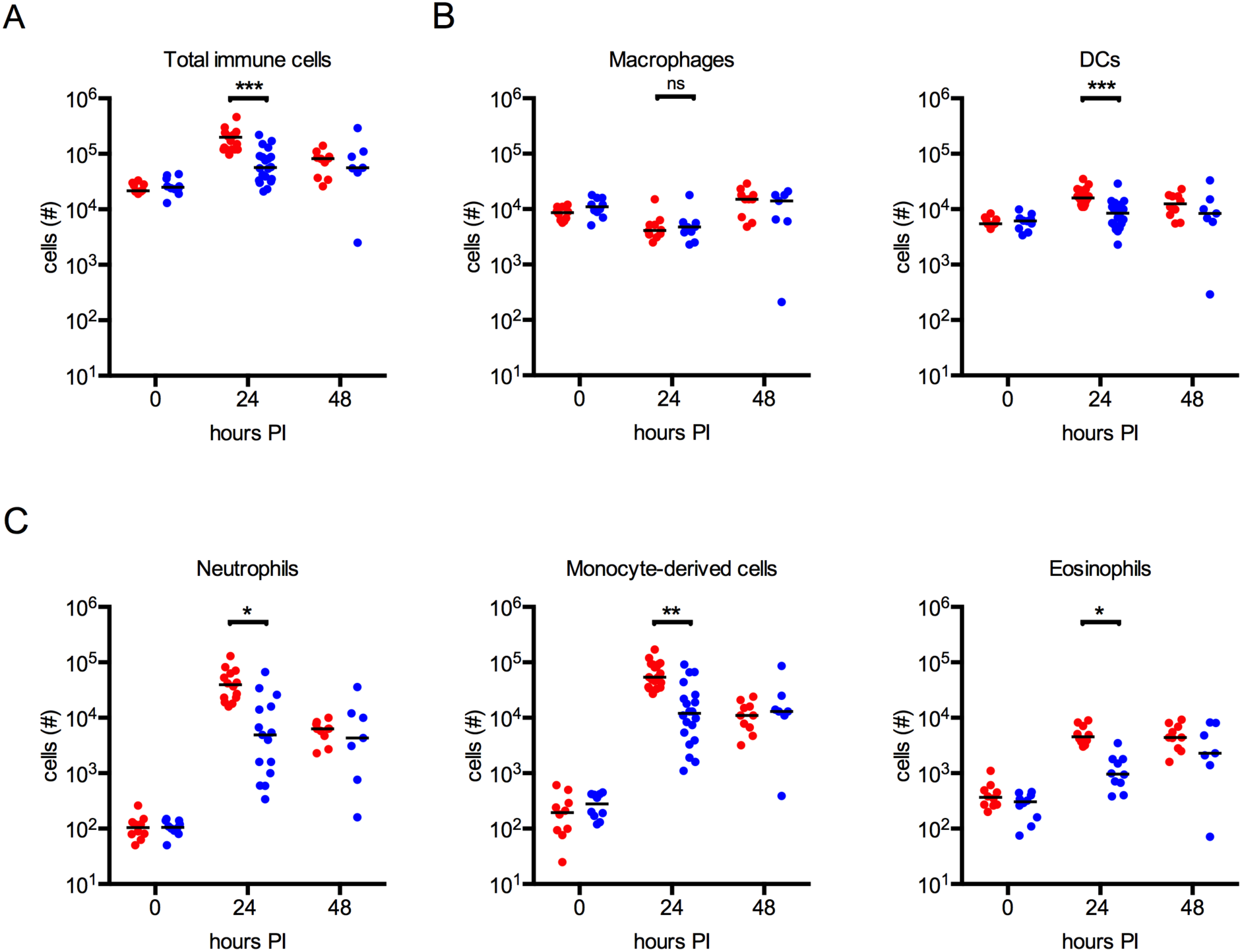
Immune cell infiltration is greater in female mice than male animals following UPEC infection. Female and male mice were infected with 10^7^ CFU of UPEC UTI89-RFP-kan^R^ for 24 or 48 hours and bladders analyzed by flow cytometry. **Figure S2** depicts gating strategies. Graphs depict (**A**) total CD45^+^ immune cells and (**B**-**C**) total specified immune cell populations at the depicted hours post-infection (PI) in bladders, 0 hours PI = naïve mice. Data are pooled from 2-3 experiments, n=3-7 mice per experiment. Each dot represents one mouse, red denotes female mice and blue indicates male mice, lines are medians. * p<0.05, ** p<0.01, *** p<0.001, ns = not significant, Kruskal-Wallis test comparing female to male at each timepoint, with Dunn’s post-test to correct for multiple comparisons.

### Bacterial uptake is altered in male mice

Previously, we reported that bacteria are found predominantly in resident macrophages at 4 hours PI and are distributed among all phagocytic cells at 24 hours PI in female mice (Mora-Bau et al., 2015). As neutrophils and monocyte-derived cells phagocytose significant numbers of bacteria in female mice (Mora-Bau et al., 2015), we reasoned that the reduced immune cell infiltration in male mice would lead to altered bacterial uptake. To test this hypothesis, we infected female and male mice with UPEC strain UTI89-RFP-kan^R^ and analyzed bladders by flow cytometry 24 hours PI to assess the identity and the number of phagocytes containing bacteria. To have a global view of UPEC-infected cells, we overlaid traditional gated populations on a tSNE plot of concatenated infected bladder samples (**Fig. 4A-B**). We then visualized UPEC-containing immune cells using the same tSNE-generated parameters in female or male bladders, observing that overall, UPEC distribution was similar, but female mice had more infected cells compared to male mice (**Fig. 4C**), which was surprising given that similar numbers of UPEC CFU were present in male and female bladder tissue at 24 hours PI (**Fig. 1B**). We quantified the number of infected cells and observed that the majority of UPEC^+^ immune cells were neutrophils and monocyte-derived cells in both sexes at 24 hours PI (**Fig. S2C-D**, gating strategy, **Fig. 4D**, MHC II^+^ and MHC II^−^ monocyte-derived cells are combined in one graph). Compared to female mice, however, male animals had fewer UPEC^+^ monocyte-derived cells, UPEC^+^ resident macrophages, and UPEC^+^ DCs at 24 hours PI (**Fig. 4D**). To test whether differences in the number of UPEC^+^ cells was a result of differential phagocytic capacity between the sexes, we assessed the percentage of UPEC^+^ cells among defined immune cell populations. While equivalent percentages of neutrophils and monocyte-derived cells contained UPEC in both sexes, female mice had a greater percentage of UPEC^+^ resident macrophages and DCs compared to male mice, suggesting that the phagocytic capacity of professional APCs may be different between the sexes (**Fig. 4E**).

**Figure 4.**
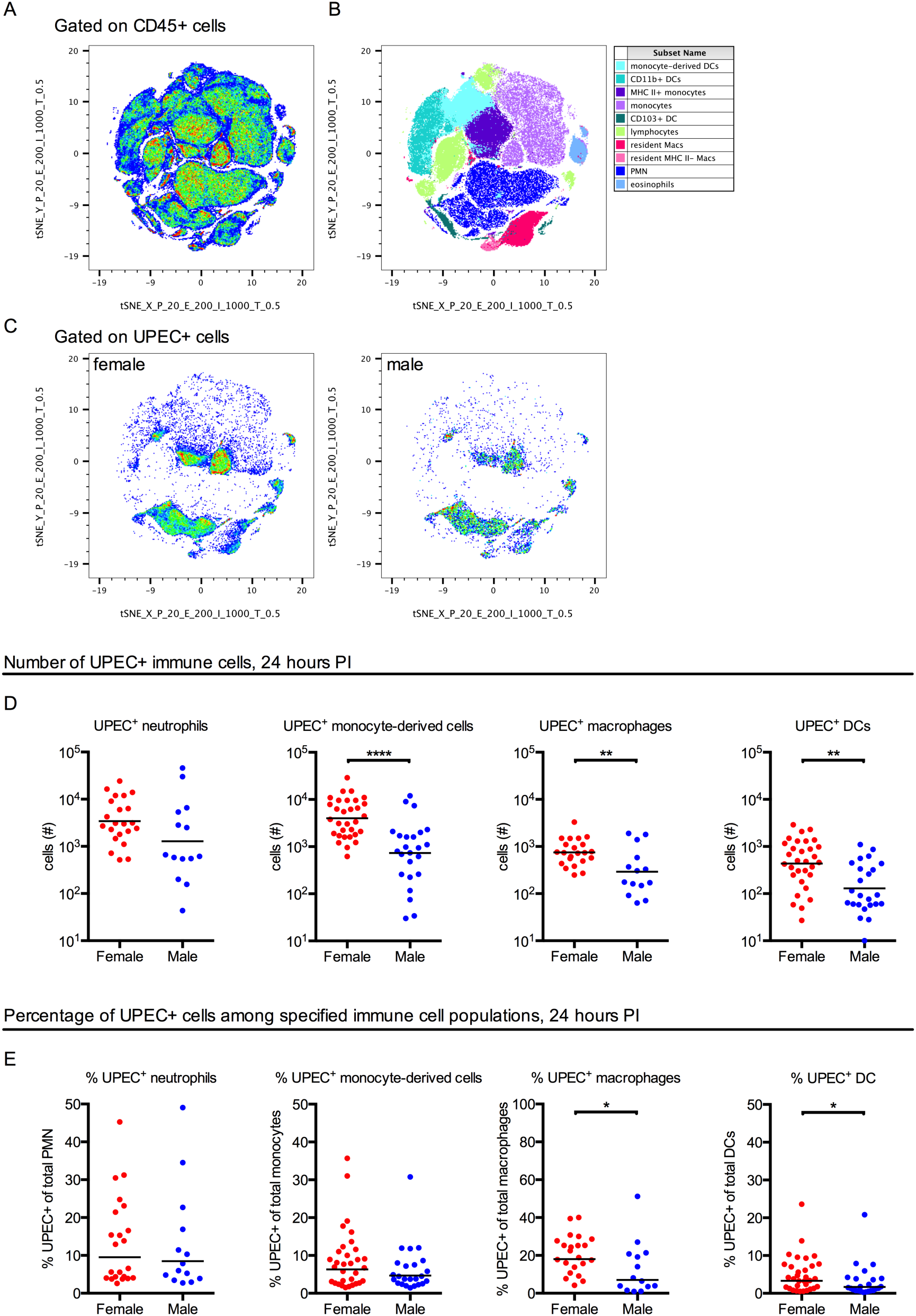
Immune cell populations contain more bacteria in female mice. Female and male C57BL/6J mice were infected with 10^7^ CFU UPEC strain UTI89-RFP-kan^R^ and bladders analyzed by flow cytometry. **Figure S2** depicts gating strategies. (**A-B**) tSNE plots show total CD45^+^ immune cell populations at 24 hours PI from (**A**) a concatenated sample of 4 female and 4 male mice and (**B**) the same plot with conventionally gated populations overlaid. (**C**) UPEC^+^ cells from female (left) or male (right) are shown using the same tSNE parameters in **A-B**. (**D**-**E**) Graphs show (**D**) the total number of specified immune cell populations containing UPEC and (**E**) the percentage of UPEC^+^ cells within the specified immune cell population. Data are pooled from 2-4 experiments, n=4-5 mice per experiment. Each dot is one mouse, red dots depict female mice and blue dots are male mice, lines are medians. * p<0.05, ** p<0.01, **** p<0.0001, Mann-Whitney test. Analyses in this figure were corrected for multiple testing by Holm– Bonferroni method, all p<0.05 had q< 0.05.

### Testosterone abrogates immunity to UTI in female mice

As hormones are major drivers of sexual dimorphism, we hypothesized that testosterone was responsible for the observed sex difference in bacterial clearance. To specifically test this, we castrated or sham-castrated 7 week old male mice to eliminate the primary source of testosterone, and allowed a 1 week recovery period before infection with UPEC. Resolution of infection was measured by monitoring for the presence of bacteria in urine, and at 28 days PI, bacterial titers were measured in the bladder. We observed no significant differences in bacterial clearance over time or in bacterial burden between mock-treated or castrated mice (**Fig. 5A-B**) although testosterone levels were efficiently decreased in castrated male mice (**Fig. 5C**).

**Figure 5.**
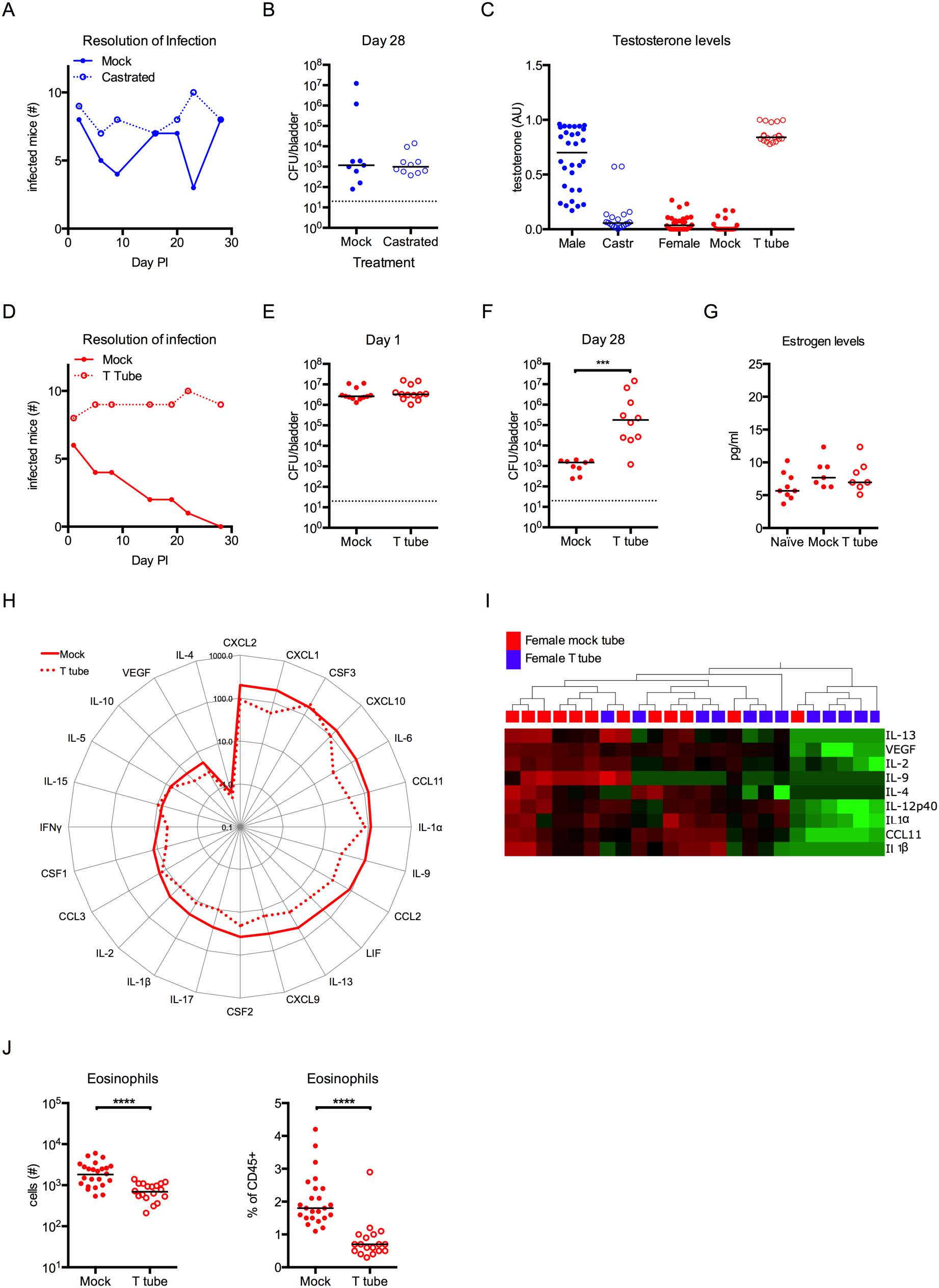
Testosterone treatment induces persistent infection in female mice. (**A-C**) 7-week old male C57BL/6J mice were castrated or mock-castrated (Mock) and allowed to recover 1 week before infection. (**C-J**) Female mice were implanted with slow release tubing containing testosterone (T tube) or empty tubing (Mock) and allowed to recover 1 week before infection. All mice were infected with 10^7^ CFU UPEC strain UTI89-GFP-amp^R^ or UTI89-RFP-kan^R^. Graphs show (**A**) the number of infected mice over time in a representative experiment, (**B**) CFU/bladder at 28 days PI in male mice, (**C**) testosterone levels in naive and castrated (Castr) male mice, and naive female mice, empty tube (Mock), and testosterone-treated (T tube) female mice, (**D**) the number of infected mice over time, (**E**) CFU/bladder at 24 hours PI, and (**F**) CFU/bladder at 28 days PI, (**G**) estrogen levels in naive female mice, empty tube (Mock), and testosterone-treated (T tube) female mice. (**H**) Spider plot shows cytokine expression 24 hours PI in mock and testosterone-treated female mice. (**I**) Heat map shows unsupervised hierarchical clustering of the 9 significantly different analytes between mock- and testosterone-treated female mice at 24 hours PI, analyzed by T-test, false discovery rate adjusted p value; q<0.15. Analytes, p, and q values are listed in **Table 2**. (**J**) Graphs show the total number and percentage of eosinophils in bladders 24 hours PI. See **Figure S3** for additional immune cell populations. Data are pooled from 2-4 experiments, n=4-6 mice/group in each experiment, except in **A** and **E**, which are a single experiments representative of 2-4 independent experiments. In **B**, **C**, **D**, **F, G, J** each dot is one mouse, lines are medians. In **F**, **I**, *** p<0.001, **** p<0.0001, Mann-Whitney test, FDR q=0.0009 to correct for multiple testing of immune populations analyzed in this figure and accompanying **Figure S3**.

We next tested the impact of testosterone in female mice using slow-release silastic tubes filled with testosterone or left empty as a control. Animals were permitted to recover 1 week after subcutaneous implantation of tubes before infection with UPEC and bacteriuria was followed over 28 days. Testosterone implantation uniformly elevated plasma testosterone levels in female mice to within the range observed in male mice (**Fig. 5C**). Strikingly, testosterone-treated female mice failed to resolve infection and had persistent bacteriuria for the duration of the experiment, whereas all mock-treated animals had sterile urine by 28 days PI (**Fig. 5D**). Additionally, although testosterone treatment did not alter bacterial burden at 24 hours PI between the two groups (**Fig. 5E**), testosterone-treated mice had significantly elevated bladder bacterial burden compared to control mice at 28 days PI, similar to that observed in male mice (**Fig. 5F**). As testosterone can be aromatized to estrogen (Simpson and Davis, 2001), we measured serum levels of estrogen in testosterone-treated female mice, observing that estrogen levels were not elevated above levels in naive or mock-tube implanted female mice (**Fig. 5G**). We assessed cytokine expression at 24 hours PI, observing that, similar to infected male mice, testosterone-treated female mice appeared to have lower overall cytokine expression levels compared to control female animals implanted with empty tubing (**Fig. 5H**). Due to the lower number of samples (n=12/group), which limited the power in this study, statistical analysis and correction for multiple testing revealed that 9/26 analytes were significantly differently expressed (q<0.15, **Fig. 5I**, **Table 2**) between the two groups. CCL11 (eotaxin), was among these cytokines, and accordingly, diminished expression correlated with a decrease in eosinophil infiltration in testosterone-treated mice (**Fig. 5J**). No differences in cellular infiltration were observed for any other cell type between mock-treated and testosterone-treated female mice 24 hours PI (**Fig. S3A**). Finally, we were surprised to observe that bacterial uptake, by any cell type, was not altered in persistently infected, testosterone-treated female mice at 24 hours PI (**Fig. S3B**), suggesting that chronic infection is not directly mediated by reduced phagocytosis early in infection.

**Table 2:**
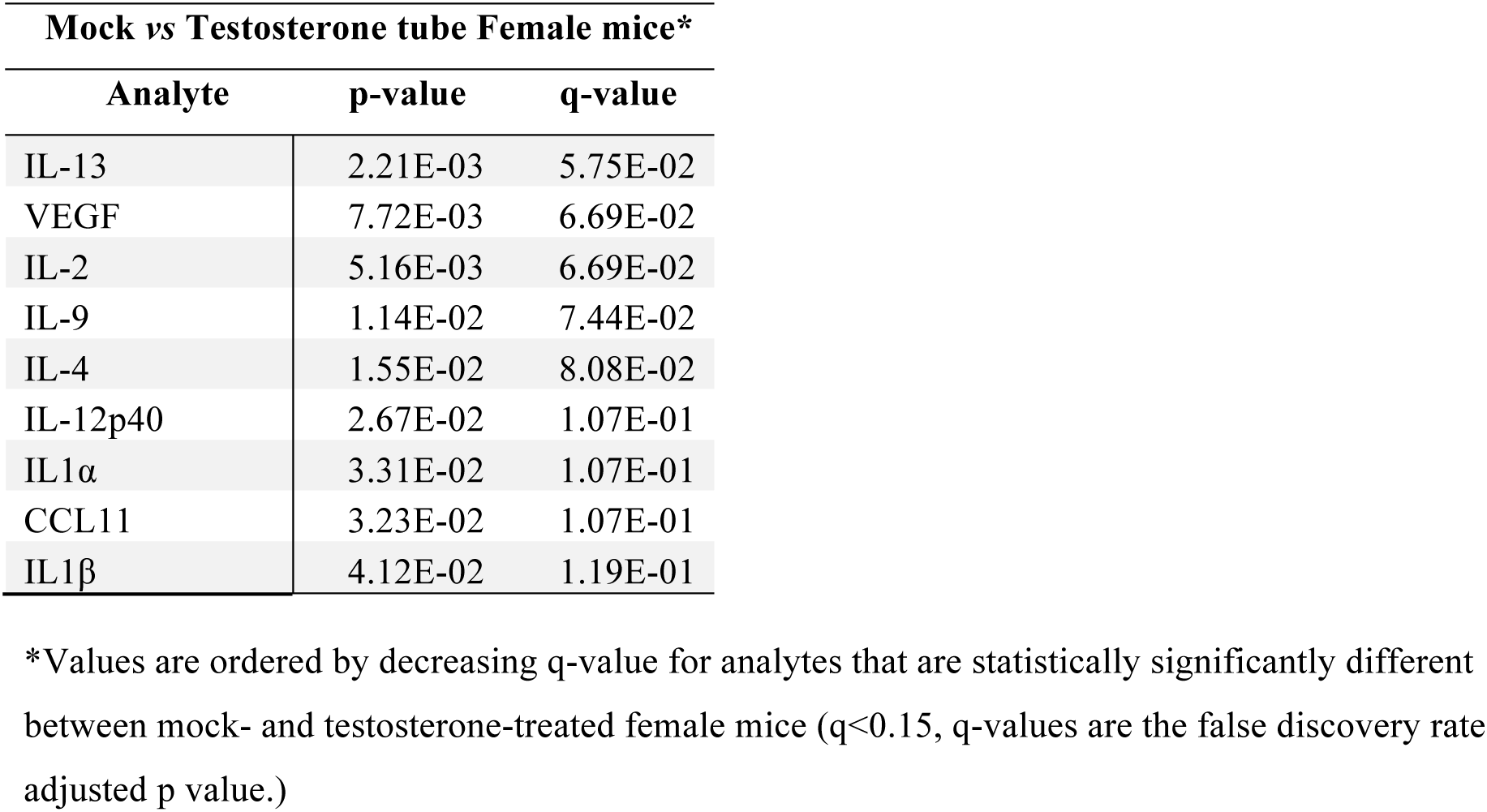
Analytes defining the response to infection in female mice treated with mock tubes compared to testosterone tubes at 24 hours.

### Female mice develop an immune response characteristic of type II immunity

IL-4, IL-5, and IL-13 were significantly elevated in female mice compared to male mice during infection and IL-4, IL-9, and IL-13 were more highly expressed in control female mice compared to those treated with testosterone (**Fig. 2** and **Fig. 5**). In addition, female mice had greater numbers of eosinophils infiltrating infected tissue compared to male mice and testosterone-treated female mice (**Fig. 5J**). Therefore, we hypothesized that female mice initiate a type II immune response mediating resolution of infection that is not induced in male mice. We first tested whether ablation of eosinophils, by administration of α-SiglecF antibody (Zimmermann et al., 2008), impacted resolution of infection. Despite reducing the number of circulating eosinophils by 60-70% (**Fig. S4A**), depleting antibody treatment did not alter the kinetic of resolution in female or male animals over 28 days (**Fig. S4B**) or the development of an adaptive immune response (**Fig. S4C**).

We then tested upstream mediators of type II immunity. Type II immunity can be initiated by damage to an epithelial cell layer, which is a hallmark of UTI (Mulvey et al., 1998). Damage results in the release of alarmins, including TSLP, IL-25, and IL-33, which induce IL-4, IL-5, IL-9, and IL-13 expression, alternative macrophage activation, and innate lymphoid cell (ILC) activation or expansion (Divekar and Kita, 2015; Fort et al., 2001; Walker and McKenzie, 2013). We measured IL-33 and IL-25 expression in bladder tissue finding that, remarkably, IL-33 expression was nearly three times higher in female mice compared to male mice at 24 hours PI (**Fig. 6A**). IL-25 was not detected in naïve or infected bladder tissue from either sex. Bladder resident macrophages from female mice upregulated IL-4Rα, which is associated with alternative activation (Gordon and Martinez, 2010), to a greater extent than in male mice during infection (**Fig. 6B**). Additionally, the number of bladder-associated ILCs, defined as CD90^+^ CD25^+^ CD3^−^ CD4^−^ NK1.1^−^ MHC II^−^ CD11b^−^, increased only in female mice 24 hours PI (**Fig. 6C**).

**Figure 6.**
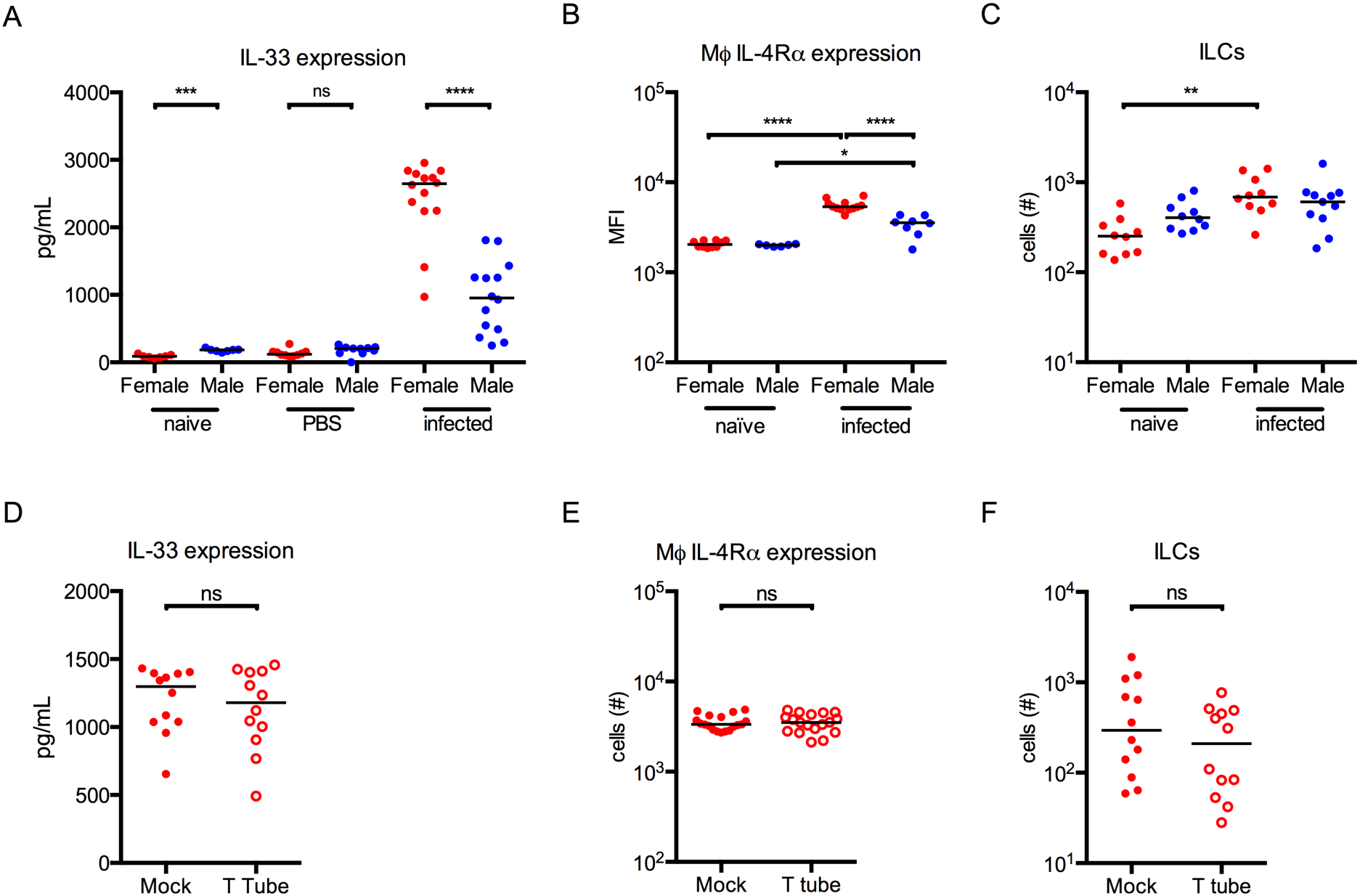
Female mice display characteristics of a type II immune response that is not suppressed by testosterone. (**A-C**) Female and male mice were infected with 10^7^ CFU of UPEC UTI89-RFP-kan^R^ and bladders analyzed at 24 hours PI. Graphs show (**A**) IL-33 protein levels in homogenized bladder, (**B**) IL-4Rα expression on bladder resident macrophages, and (**C**) the number of ILCs (CD90^+^ CD25^+^ CD3^−^ CD4^−^ NK1.1^−^ MHC II^−^ CD11b^−^) in bladders. (**D-F**) Female mice were implanted with empty tubing (Mock) or slow release tubing containing testosterone (T tube) and allowed to recover 1 week before infection with 10^7^ CFU UPEC strain UTI89-RFP-kan^R^. Graphs show (**D**) IL-33 protein levels in homogenized bladder tissue, (**E**) IL-4Rα expression on bladder resident macrophages, and (**F**) the number of ILCs in bladders. Data are pooled from 2-3 experiments, n=4-5 mice/group in each experiment. Each dot is one mouse, red dots depict female mice and blue dots are male mice, lines are medians. ns = not significant, ** p<0.01, *** p<0.001, **** p<0.0001 Mann-Whitney test. Analyses in this figure were corrected for multiple testing by Holm–Bonferroni method, all p<0.05 had q< 0.05.

We reasoned that exogenous IL-33 would promote resolution of infection in male mice. Male mice were infected and IL-33 was administered directly into the bladder at the time of infection and 24 hours PI (**Fig. S5A**). No differences in resolution were observed between the treated and control groups (**Fig. S5B**). In addition, IL-33 supplementation did not impact bacterial burden between treated and control animals following primary or challenge infection (**Fig. S5C**). We also tested whether neutralizing IL-33 in female mice would impair bacterial clearance or innate immune responses (**Fig. S5A**) (Kim et al., 2012). Female mice, treated with an α-IL-33 antibody 2 hours prior to infection and 1, 4 and 7 days PI, resolved their infection with similar kinetics and had no differences in bacterial burden following primary or challenge infection compared to isotype-treated mice (**Fig. S5D-E**). At 24 hours PI, treated and control female mice showed no differences in the number of resident immune cells, infiltrating myeloid cells, or lymphocytes as compared to isotype-treated mice (**Fig. S5F**). The number of UPEC^+^ immune cells was also not different between the groups (**Fig. S5G**). Supporting that type II immune responses were not directly responsible for bacterial clearance, we observed no differences between control and testosterone-treated mice with respect to IL-33 expression, macrophage IL-4Rα expression, or number of ILCs in infected bladder tissue 24 hours PI (**Fig. 6D-F**), despite that testosterone treatment abrogated the capacity to resolve infection in female mice.

### IL-17 is necessary for resolution of UTI

Having ruled out a role for type II immunity in bacteria clearance, we considered whether other immune cell populations contributed to resolution. In the course of our investigations, we also quantified lymphocytic populations accumulating in the bladder during UTI. We observed that CD3^+^CD4^+^ T cells, NK cells, γδ T cells, and ILC3 LTi-like cells (CD90^+^ CD25^+^ CD4^+^ CD3^−^ NK1.1^−^ MHC II^−^ CD11b^−^) accumulated in greater numbers in female mice as compared to male mice 24 hours PI (**Fig. 7A**). Consistent with this observation, these cell populations were decreased in testosterone-treated female mice compared to control-treated mice (**Fig. 7B**). Our previous work ruled out a role for T cells in bacterial clearance (Mora-Bau et al., 2015), therefore, we infected female wildtype and *Rag2*^−/−^γc mice, which lack NK cells and LTi-like cells, and followed resolution of infection over time. Remarkably, *Rag2*^−/−^γc mice remained infected up to 80 days PI, whereas wildtype female mice resolved their infection within 1 month (**Fig. 7C**). Of note, *Rag2*^−/−^γc mice have eosinophils, supporting our conclusion that these cells do not mediate resolution of infection. To specifically test the contribution of NK cells, we infected wildtype female mice treated with a depleting NK1.1 antibody. In contrast to the outcome in infected *Rag2*^−/−^γc mice, NK cell depletion did not impact bacterial clearance and both groups of mice cleared their infection by day 28 (**Fig.7D**).

**Figure 7.**
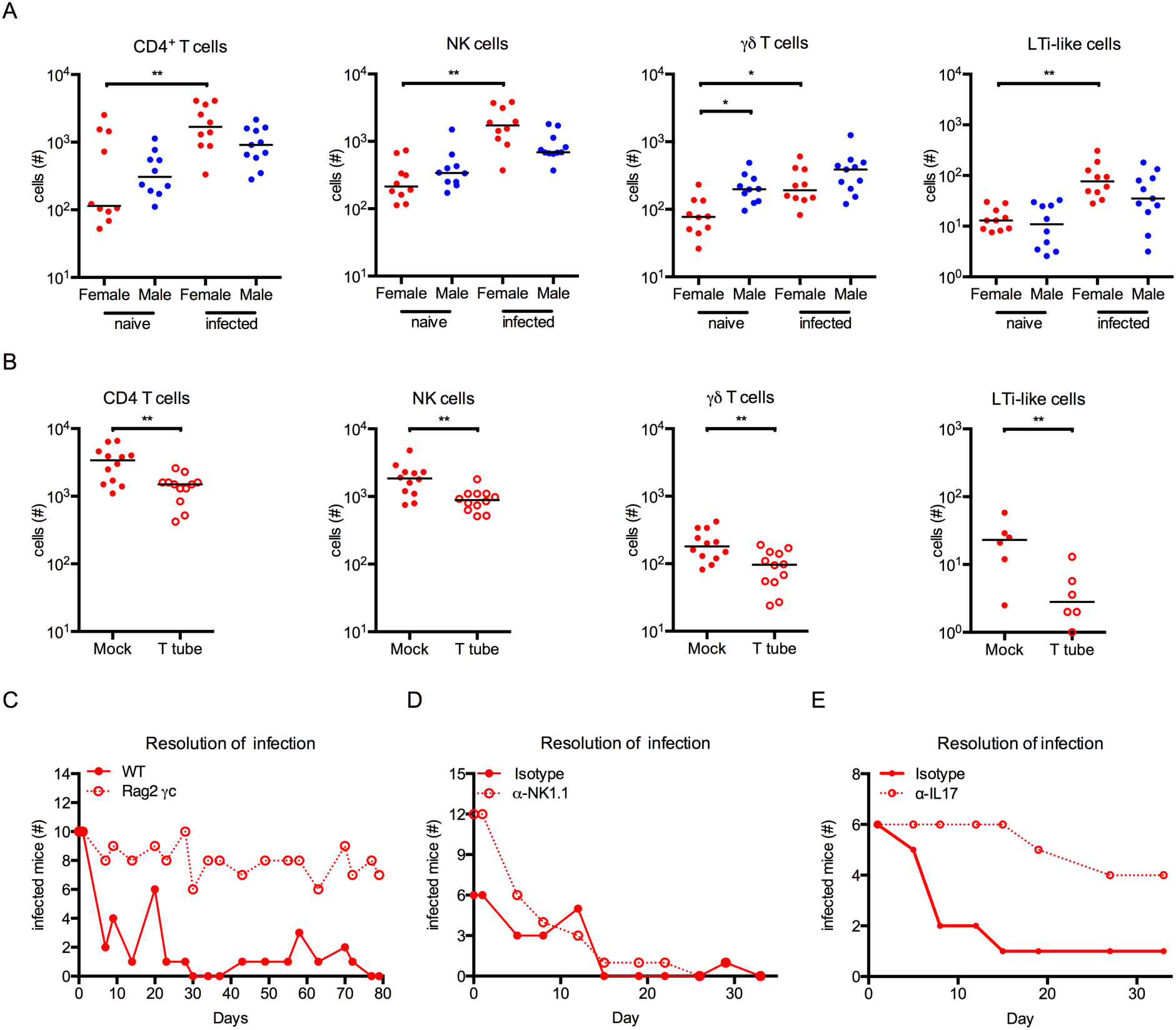
IL-17 is necessary for resolution of infection in female mice. (**A**) Female and male mice were infected with 10^7^ CFU of UPEC strain UTI89-RFP-kan^R^ and bladders analyzed by flow cytometry at 24 hours PI. Graphs show (**A**) the number of CD3^+^ CD4^+^ T cells, CD3^−^ NK1.1^+^ NK cells, γδ T cells, and LTi-like cells (CD90^+^ CD25^+^ CD4^+^ CD3^−^ NK1.1^−^ MHC II^−^ CD11b^−^) in bladders. (**B**) Female mice were implanted with empty tubing (Mock) or slow release tubing containing testosterone (T tube) and allowed to recover 1 week before infection with 10^7^ CFU UPEC strain UTI89-RFP-kan^R^. Graphs show the number of CD3^+^ CD4^+^ T cells, CD3^−^ NK1.1^+^ NK cells, γδ T cells, and CD90^+^ CD25^+^ CD4^+^ CD3^−^ NK1.1^−^ MHC II^−^ CD11b^−^ LTi-like cells in bladders. (**C-E**) Graphs show the number of infected mice, determined by urine sampling, over time in (**C**) wildtype or *RAG2^−^*/-γc mice, (**D**) mice treated with isotype or αNK1.1-depleting antibody, (**E**) mice treated with α-IL-17 neutralizing antibody. (**A-B**) Data are pooled from 2-3 experiments, n=4-5/group in each experiment. Each dot is one mouse, red dots depict female mice and blue dots are male mice, lines are medians. (**C-E**) Graphs are representative experiments, n=6-12 mice per experiment, at least 2 experiments each. * p<0.05, ** p<0.01, Mann-Whitney test. Analyses in this figure were corrected for multiple testing by Holm–Bonferroni method, all p<0.05 had q< 0.05.

Our experiments ruled out a role for NK cells in resolution, however the contribution of LTi-like cells to UTI immunity was still unclear. Interestingly, *Rag2*^−/−^ mice can express IL-17 from LTi-like cells following microbial stimulation, whereas *Rag2*^−/−^γc animals do not as they lack this cell population (Takatori et al., 2009). Of the cytokines expressed in female and male mice during UTI, IL-17 exhibited one of the largest differences between the sexes, which persisted at 48 hours PI (**Fig. 2B-C**). Thus, we hypothesized that IL-17 expression, early in infection, mediates bacterial clearance. We administered a single dose of α-IL-17 neutralizing antibody intravenously immediately prior to infecting mice with one of our two isogenic UPEC strains and monitored bacteriuria over time. Notably, 8/12 mice treated with the α-IL-17 antibody remained infected over 28-35 days, whereas 11/12 isotype-treated animals cleared bacteria with the expected kinetic (**Fig. 7E**), supporting that this cytokine pathway drives resolution of UTI.

## Discussion

Women experience UTI at a much higher frequency than men, although men develop more severe, chronic infections (Fabbian et al., 2015; Lipsky, 1989). While anatomical differences are often cited as an explanation for the disproportional incidence in UTI between the sexes, this fails to take into account that frequency of infection is equivalent between older women and men, and in infants of both sexes (Ruben et al., 1995), and does not address disparities in infection outcome in either sex. Hypothesizing that sex-based differences in immunity influence response to UTI, we directly compared the innate and adaptive immune response in infected female and male mice. Female mice displayed a superior capacity to resolve infection compared to male mice and testosterone-treated female mice. Remarkably, IL-17 neutralization induced chronic infection in female mice.

IL-17 initiates many anti-bacterial pathways, including antimicrobial peptide and chemokine expression, recruitment and activation of neutrophils and monocytes, and potentially direct killing of infected cells (Kolls et al., 2008; Papotto et al., 2017; Veldhoen, 2017). Supporting that IL-17 contributes to UTI resolution, bladder-associated bacterial CFU are higher in IL-17a^−/−^ mice compared to wildtype animals at 3 and 4 days PI, although whether the mice develop chronic infection is unknown (Sivick et al., 2010). T_h_17 T cells, γδ T cells, neutrophils, and LTi-like cells express IL-17 (Papotto et al., 2017; Takatori et al., 2009). Typically, γδ T cells express high levels of IL-17 following infection or in the context of autoimmune inflammation (Papotto et al., 2017). While γδ T cells are absent in *Rag2*^−/−^ mice, LTi-like cells are present (Takatori et al., 2009), which may explain why female *Rag2*^−/−^ mice resolve UTI with wildtype kinetics (Mora-Bau et al., 2015), whereas *Rag2*^−/−^γc animals remained chronically infected in this study.

IL-17-mediated inflammation displays sex differences. In an allergic asthma model, male mice exhibit less inflammation compared to female mice and administration of testosterone leads to decreased numbers of IL-17-expressing T_h_17 T cells and an overall reduction in the level of inflammation associated with lung injury (Fuseini et al., 2018). Perhaps, most intriguingly, genetic variation in the Y chromosome imparts varying degrees of protection in a model of flu infection. In this study, infection of Y chromosome consomic C57BL/6 mice revealed that Y chromosome polymorphisms are associated with differences in the frequency of γδ T cells and IL-17 expression, which subsequently impact inflammatory response and disease severity in the lungs (Krementsov et al., 2017).

IL-17 may also drive the initiation of pro-resolving type II immunity in the bladder. Indeed, in adipose tissue, the absence of γδ T cells or IL-17 leads to a reduction in IL-33 (Kohlgruber et al., 2018). In this tissue, TNFα and IL-17 induce upregulation of IL-33, which in turn, controls thermoregulation and response to cold shock (Kohlgruber et al., 2018). Notably, TNFα is highly expressed in the bladder in the first hour following UPEC infection and, while IL-17 peaks at 24 hours PI, its expression is increased 10-100 fold over uninfected female animals 1-6 hours PI (Ingersoll et al., 2008), potentially inducing the increase in IL-33 observed in this study. The presence of a type II immune signature was unexpected. Type II immunity is typically associated with parasitic infections and non-infectious inflammatory diseases such as asthma and very few reports have focused on bacterial infection (Divekar and Kita, 2015; Gieseck et al., 2018; Hogan et al., 2013). However, the bladder retains metabolic waste until excretion and thus, repair of infected or damaged tissue must occur rapidly. Thus, we speculate that this signaling cascade acts to quickly induce type II immunity pathways to restore the important barrier function of the bladder. While this concept remains to be tested, emerging evidence supports that IL-17 and IL-33 work in concert to maintain tissue homeostasis (Kohlgruber et al., 2018).

The cytokine profile observed in female mice suggests that ILCs may play a role in the induction of inflammation (LTi-like cells or ILC3s) or reparative functions (ILC2s) following infection. Notably, in the lung, ILCs, in female but not male mice, responded to IL-33 treatment by production of large amounts of IL-5 and IL-13, and cytokine response can be dampened by testosterone treatment (Cephus et al., 2017; Warren et al., 2017), suggesting that sex differences in the biology of ILCs influence immunity. This is an exciting new horizon for bladder mucosal immunity as, to the best of our knowledge, bladder-resident ILCs have not been reported. A single publication has described the presence of ILC2s in the urine of patients undergoing BCG immunotherapy for bladder cancer, however, their origin and function remain unknown (Chevalier et al., 2017). Activation and expansion of macrophages and ILCs following IL-33 expression would be expected to induce or amplify expression of IL-4, −5, −9, and −13, as well as CCL11, mediating eosinophil infiltration (Chen et al., 2004; Heller et al., 2012; Mould et al., 1997). Thus, eosinophils, while not required for bacterial clearance, likely play a role in another aspect of host defense, repair, or homeostasis. While this is an area that clearly requires further investigation, one possibility is that eosinophils amplify the type II immune response supporting humoral responses following infection.

What remains to be determined is the underlying cause of this differential cytokine response between female and male mice. Development of chronic UTI in male and testosterone-treated female mice supports that sex hormone levels play an essential role in the response to bacterial colonization. Although we ruled out that testosterone was aromatized to estrogen in our study, estrogen can also play a direct or indirect role in susceptibility to infection, particularly in older patients. Signaling via estrogen receptor α enhances IL-4 and IL-13 expression and polarization of myeloid cells towards an M2-like phenotype (Campbell et al., 2014; Keselman and Heller, 2015). Estrogen treatment *in vitro* increases human β-defensin 3 expression (Luthje et al., 2013), although how this is mediated is unknown. Ovariectomized mice have higher bladder bacterial colonization compared to intact mice (Luthje et al., 2013), however, this was not recapitulated in the surgical model comparing female and male UTI (Olson et al., 2016). One reason may be that, in addition to the method employed to infect mice (*i.e.*, catheterization of female mice *versus* incision through the abdominal wall in both sexes), the mouse and bacterial strains differ between these two studies (Luthje et al., 2013; Olson et al., 2015). Notably, our study utilized one of the same mouse strains and the same bacterial strain as those used in Olsen *et al.*, however, C57BL/6J mice remained chronically infected only in our study, suggesting that response to surgical manipulation may impact bacterial clearance (Olson et al., 2015).

In sum, our study demonstrates, in a physiologically relevant model that IL-17 specifically influences the innate response to bacterial infection in a sex-dependent manner. Interestingly, the first 24 hours of infection were critical in determining whether an animal would resolve infection. Whether the response to infection in this same time period in humans determines outcome remains to be seen, however our analysis of human samples revealed that a significant number of sex differences in immune gene expression, including players in the IL-17 - IL-33 axis, arise in the first 22 hours following *E. coli*-induced immune responses in whole blood. Thus, our work may unlock the potential for fundamental research to better understand immunity in this underappreciated organ, by providing a valuable tool for the investigation of urogenital pathologies that display a sex difference between men and women beyond UTI, such as bladder cancer or interstitial cystitis, and support the development of sex-specific immune modulating therapies to improve the pronounced sex differences observed clinically in response to infection of the urinary tract.

## Materials and Methods

### Study Design

This study was conducted using a preclinical mouse model in controlled laboratory experiments to test the hypothesis that the immune response to urinary tract infection differs between female and male animals. Animals were assigned to groups upon random partition into cages. In all experiments, a minimum of 3 and a maximum of 12 animals made up an experimental group and all experiments were repeated 2-5 times. Data were pooled before statistical analysis and outliers were identified using the Grubbs’ test (GraphPad QuickCalcs). In the case of an outlier in a parameter, all parameters measured in the corresponding mouse were excluded. As determined *a priori*, all animals having abnormal kidneys (atrophied, enlarged, white in color) at the time of sacrifice were excluded from all analyses, as previously we observed that abnormal kidneys negatively impact resolution of infection. Endpoints were determined prior to the start of experiments and researchers were not blinded to experimental groups.

### Ethics Statement

Animal experiments were conducted in accordance with approval of protocol number 2012-0024 and 2016-0010 by the *Comité d’éthique en expérimentation animale Paris Centre et Sud* and the *Comités d’Ethique pour l’Expérimentation Animale* Institut Pasteur (the ethics committee for animal experimentation), in application of the European Directive 2010/63 EU. In all experiments, mice were anesthetized by intraperitoneal injection of 100 mg/kg ketamine and 5 mg/kg xylazine and sacrificed either by cervical dislocation or carbon dioxide inhalation. Human data was from the *Milieu Intérieur* Project (https://clinicaltrials.gov; identifier: NCT01699893) and was previously described (Piasecka et al., 2018).

### Mice

Female and male between the ages of 6-12 weeks were used in this study. C57BL/6J were obtained from Charles River Laboratories France, *RAG2^−^*/-γc were bred in house at Institut Pasteur, Paris.

### Urinary tract infection and determination of bacterial burden

Female and male mice between 6 and 8 weeks of age were anesthetized as above, catheterized transurethrally, and infected with 10^7^ colony forming units of UTI89-GFP-amp^R^ or UTI89-RFP-kan^R^ (Mora-Bau et al., 2015) in 50 µL PBS as previously described (Hung et al., 2009; Zychlinsky Scharff et al., 2017). UTI89-GFP-amp^R^ and UTI89-RFP-kan^R^ infect with equal efficiency (Mora-Bau et al., 2015) and were used interchangeably, with the exception of experiments to detect bacteria by flow cytometry, in which only UTI89-RFP-kan^R^ was used because the GFP signal is masked by autofluorescence in the bladder (Mora-Bau et al., 2015).

Infection was monitored by examining bacterial growth from urine samples. Urine was collected every 2-5 days and 2 µL were diluted directly into 8 µL PBS spotted on agar plates containing antibiotics as appropriate. The presence of any bacterial growth was counted as positive for infection. The limit of detection (LOD) for this assay is 500 bacteria per mL of urine. Mice were sacrificed at indicated timepoints, bladders homogenized in sterile PBS, serially diluted, and plated to determine CFU. The LOD for CFU is 2 or 3×10^1^ bacteria per organ, depending upon the number of dilutions plated, and is indicated by a dotted line in graphs. All sterile organs are reported at the LOD.

### Flow cytometry

Mice were sacrificed at indicated timepoints and the bladders removed. Single cell homogenates were prepared as previously described (Mora-Bau et al., 2015). Briefly, minced bladders were incubated in 0.34 Units/mL Liberase TM (Roche) diluted in PBS at 37°C for 1 hour, with manual agitation every 15 minutes. Digested tissue was passed through a 100 µm filter (Miltenyi), washed, blocked with FcBlock (BD Biosciences), and immunostained (**Supplemental Table 2**). Samples were acquired on a BD Fortessa (BD Biosciences) and analyzed using FlowJo Version 10 software.

### Luminex MAP analysis

Bladders were removed at 0, 24, and 48 hours PI and placed in 1 mL of cold PBS and homogenized with a handheld tissue grinder on ice or with a PreCellys24 bead mill homogenizer. 100 µL were removed to determine CFU and homogenates were clarified by microcentrifugation (13K, 4°C, 5 minutes) and stored at −20**°**C until analysis. After thawing, prior to analysis, samples were centrifuged a second time to remove remaining cell debris. All samples were assessed together to avoid inter-assay variability by Millipore Milliplex MAP Mouse Cytokine/Chemokine Magnetic Bead Panel, Premixed 32-Plex, according to the manufacturer’s recommendations (Merck Millipore). Cytokines with values less than the minimal detectable concentration +2 standard deviations (MinDC+2SD, manufacturer’s recommendations) in both female and male mice at 24 hours PI, the peak of cytokine expression (Ingersoll et al., 2008), were removed prior to statistical analysis; these were IL-3, IL-7, IL-12p70, MIP1β, RANTES, and TNFα.

### Castration, hormone implantation, and determination of hormone levels

6-7 week old male C57BL/6J mice were anesthetized and a small incision made in the scrotum. Testes were externalized, large vessels ligated, and the testes were removed. The incision was closed using VetBond glue. Control (mock) animals received an incision, the testes were exposed, and then the incision was sealed with VetBond. Empty control and testosterone-filled implants were made according to (Ketterson et al., 1991). Briefly, silastic tubing (Dow Corning) containing 5 mM packed testosterone propionate (Sigma, Cat # 86541), approximately 0.8-1 mg, were implanted under the skin of 6-7 week old female C57BL/6J mice through a small incision. Incisions were closed using 1-2 stitches with Vicryl sutures. Control (mock) animals received empty tubing. All mice were allowed to recover for one week before further manipulation. Plasma testosterone and estrogen levels were determined by ELISA following manufacturer’s instructions (Abcam).

### Cell depletion and protein neutralization

Cell depletion and cytokine neutralization were performed as follows:

For SiglecF depletion, 10 µg/mouse α-SiglecF antibody (clone E50-2440, BD Pharmingen) or 10 µg/mouse IgG2a isotype control (clone R35-95, BD Pharmingen) were delivered intraperitoneally in 100 µL PBS (Zimmermann et al., 2008) at the time of infection, and every 7 days for 3 weeks. For NK cell depletion, 100 µg/mouse α-NK1.1 (clone PK136, BioXcell) or 100 µg/mouse IgG2a isotype control (clone C1.18.4, BioXcell) were delivered intraperitoneally in 100 µL PBS. Mice received isotype control or depleting α-NK1.1 24 hours prior to primary infection and then again on day 6, 13, and 20 post-primary infection. For IL-33 neutralization, 3.6 µg/mouse α-IL-33 (catalog number AF3626, BioTechne/R&D Systems) or 3.6 µg/mouse polyclonal goat IgG isotype control (catalog number AB-108-C, BioTechne/R&D Systems) were delivered intraperitoneally in 100 µL PBS (Kim et al., 2012). Mice received isotype control or neutralizing α-IL-33 2 hours prior to primary infection and then again on day 1, 4, and 7 post-primary infection. For IL-33 supplementation, mice received 2.5 ng recombinant IL-33 (equivalent to that measured in female mice at 24 hours PI) (catalog number 3626-ML-010, BioTechne/R&D Systems) intravesically at the time of infection and 24 hours PI. For IL-17 neutralization, 83.3-111 µg/mouse α-IL-17 (clone 50104, BioTechne/R&D Systems) or 83.3-111 µg/mouse rat IgG2a isotype control (clone 54447, BioXcell) were delivered intravenously in 100 µL PBS. Mice received a single injection of isotype control or neutralizing α-IL-17 at the time of infection.

### Statistical analysis

Statistical significance was performed in Prism 6 for Mac OS X using the nonparametric Mann-Whitney test (two groups) or the nonparametric Kruskal-Wallis test with a Dunn’s multiple comparison post-test (3 or more groups) for all data except cytokine analysis and human mRNA expression data. Multiple testing correction (*q*) within analyses was performed using the Holm–Bonferroni method (https://jboussier.shinyapps.io/MultipleTesting/). Qlucore Omics Explorer 3.2 software was used to test statistical significance of cytokine protein expression data and human mRNA expression data after log-transformation of the raw data and to calculate *q*-values (the false discovery rate adjusted *p* value) after log transformation of the values, using either a T-test or ANOVA as indicated.

## Acknowledgments

We gratefully acknowledge insightful discussions, technical support, and/or critical reading of the manuscript by Drs. Melanie Hamon, Lucy Glover, Jessica Quintin, Vincent Rouilly, Kimberly Kline, Nicolas Serafini, Clémence Hollande, and Björn Albrecht. We acknowledge the Labex Milieu Intérieur and the Technology Core of the Center for Translational Science at the Institut Pasteur for supporting aspects of this study. We thank Dr. Tim Sparwasser for sponsoring the research stay of AZS.

## Author contributions

Conceptualization: AZS, MR, LLM, and MAI; Methodology: AZS, MF, MAI; Investigation and data analysis: AZS, MR, LLM, TC, DD, MAI; Writing - Original Draft: AZS and MAI; Writing - Review & Editing: AZS, MR, LLM, TC, MLA, MF, DD, MAI; Funding Acquisition: MLA and MAI; Supervision: MAI.

## Supplementary Materials

**Figure S1.**
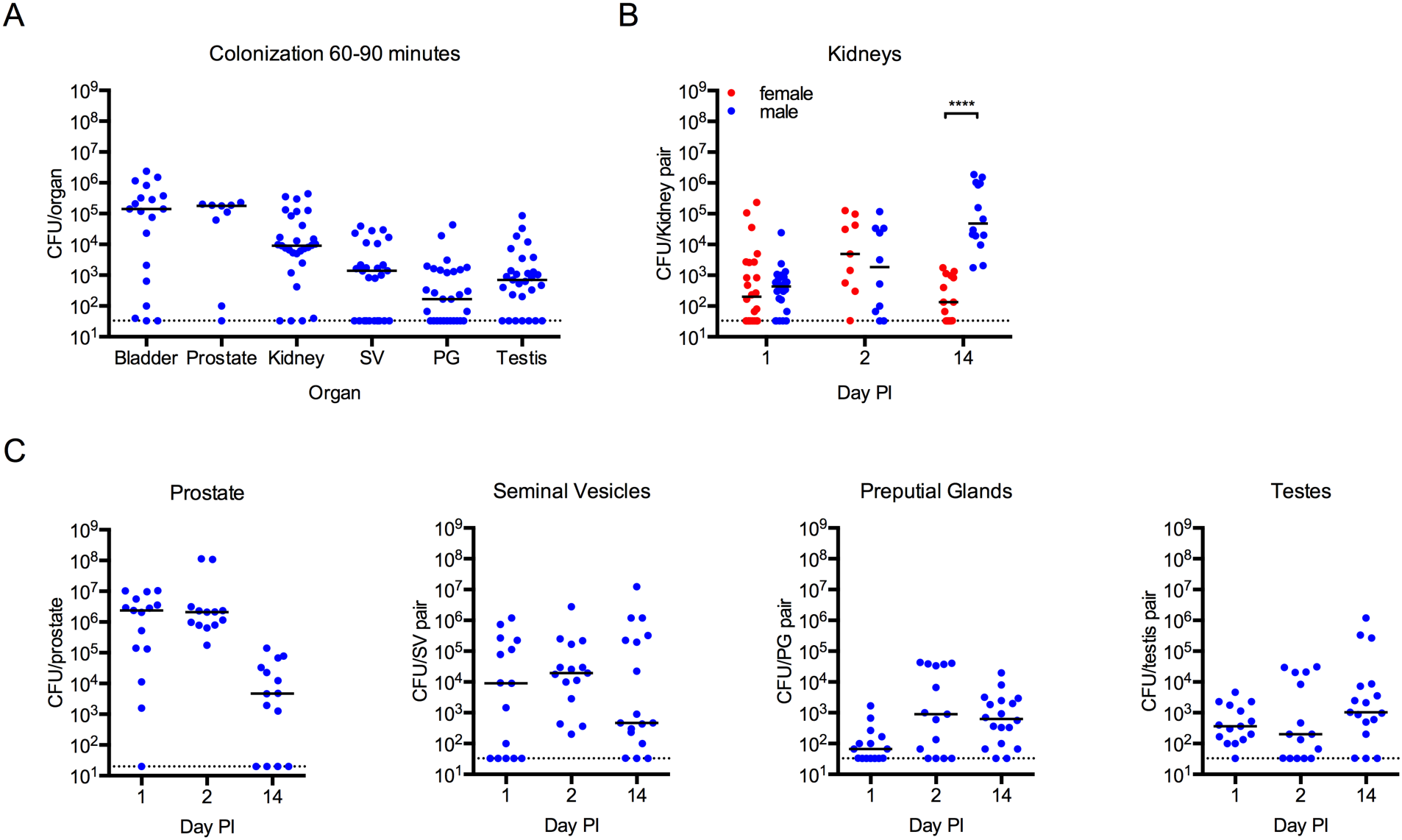
Male mice are colonized by uropathogenic *E. coli* following intravesical instillation *via* the urethra. (**A**) Male C57BL/6J mice were infected with 10^7^ colony forming units (CFU) of UPEC strain UTI89-RFP-kan^R^ and CFU/organ was determined 60-90 minutes post-infection (PI). (C) Female and male mice were infected with 10^7^ CFU of UPEC strain UTI89-GFP-amp^R^ or UTI89-RFP-kan^R^ and CFU/kidney pair was determined at the indicated times PI. (**D**) Graphs depict CFU/organ in male mice infected with 10^7^ CFU of UPEC at the indicated times PI. Data are pooled from 2-4 experiments, n=4-10 mice/group in each experiment. Each dot represents one mouse. Dotted line depicts the limit of detection for the assay, 20 or 33 CFU/organ.

**Figure S2.**
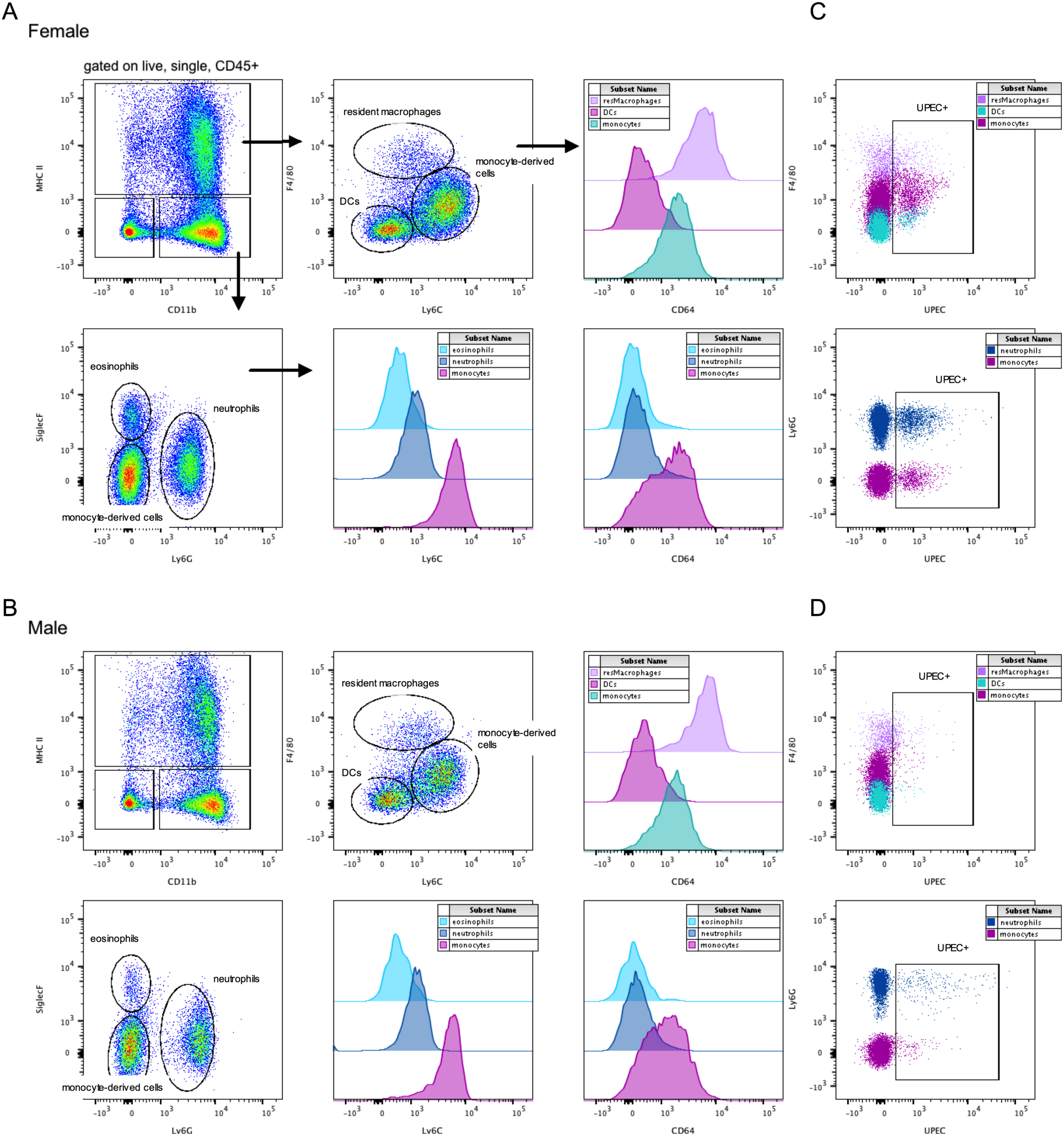
Gating strategy to identify immune cells and UPEC^+^ immune cells. Plots demonstrate the gating strategies used for analyses shown in **Figure 3**, **4**. Representative plots for (**A**) female and (**B**) male mice are shown. Following infection with 10^7^ CFU UPEC strain UTI89-RFP-kan^R^, bladders were digested and immunostained as described in the Methods and (Mora-Bau et al., 2015). Each sample, consisting of a single bladder, was acquired on a BDFortessa SORP. To identify infiltrating immune cells, samples were gated on CD45^+^ cells, excluding doublets. Resident macrophages were identified as MHC II^+^ CD11b^+^ F4/80^+^ CD64^+^Ly6C^−^ Ly6G^−^, dendritic cells were MHC II^+^ CD11b^+/−^ CD103^+/−^ CD11c^+^ F4/80^−^ CD64^−^ Ly6C^+/−^ Ly6G^−^, monocyte-derived cells were CD11b^+^ Ly6G^−^ Ly6C^+^ MHC II^−^ or MHC II^+^ F4/80^int^ CD64-/int, neutrophils were identified as CD11b^+^ Ly6G^+^ MHC II^−^ F4/80^−^ CD64^−^, eosinophils were CD11b^+^ SiglecF^+^ Ly6G^−^ MHC II^−^ F4/80^int^ CD64^−^. (**C-D**) UPEC^+^ immune cells were identified by population as specified above and then all RFP^+^ cells were gated in (**C**) female mice or (**D**) male mice.

**Figure S3.**
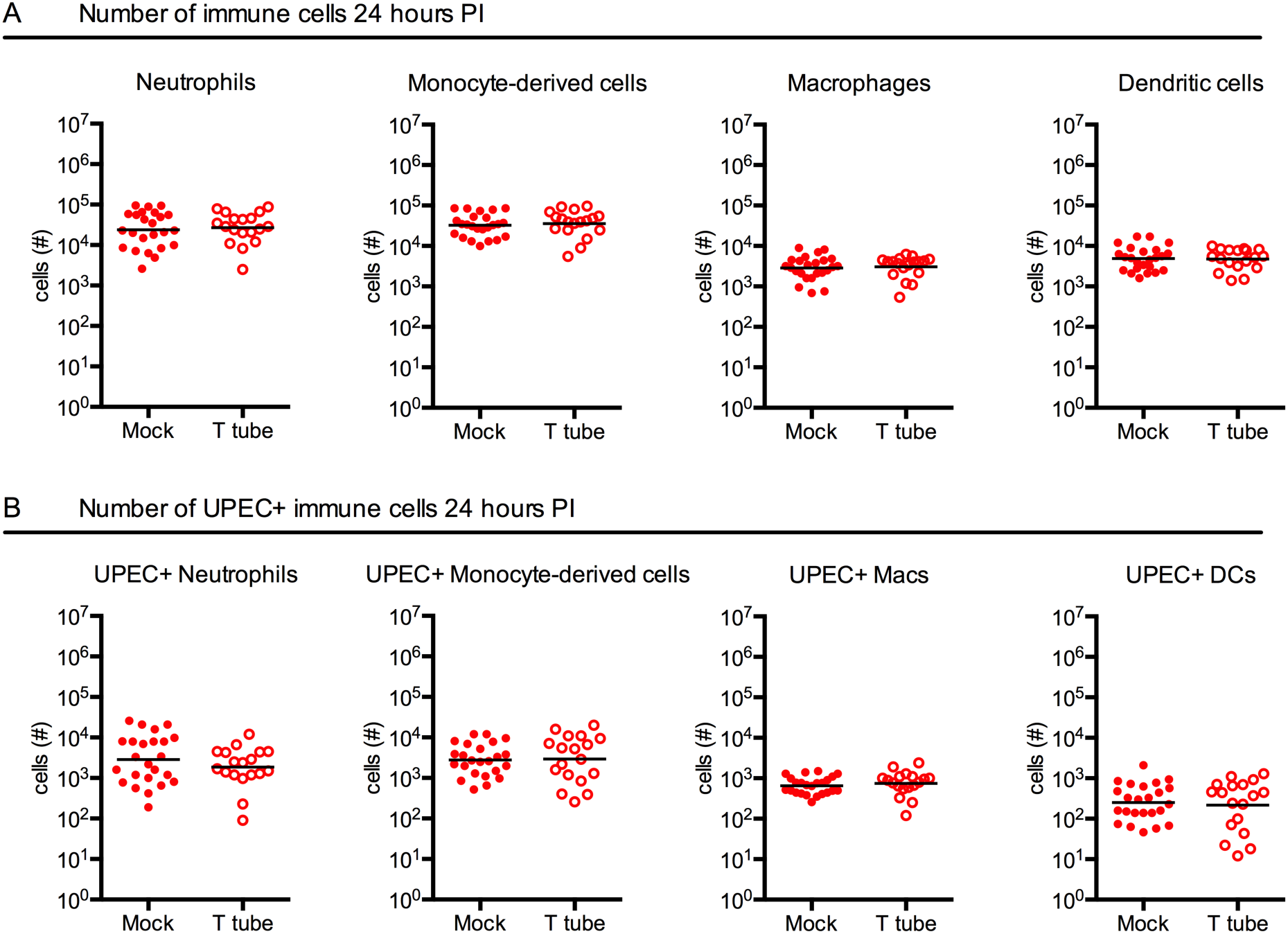
Testosterone treatment in female mice does not alter immune cell infiltration or bacterial phagocytosis. (**A**-**B**) Female C57BL/6J mice were implanted with empty tubing (Mock) or slow release tubing containing testosterone (T tube) and allowed to recover 1 week before infection with 10^7^ CFU UPEC. Graphs show the absolute number of (**A**) infiltrating immune cells and (**B**) UPEC^+^ immune cells at 24 hours PI. Data are pooled from 4 experiments, n=4-5 mice/group in each experiment, each dot is one mouse, lines are medians. Mann-Whitney test, no statistically significant differences found.

**Figure S4.**
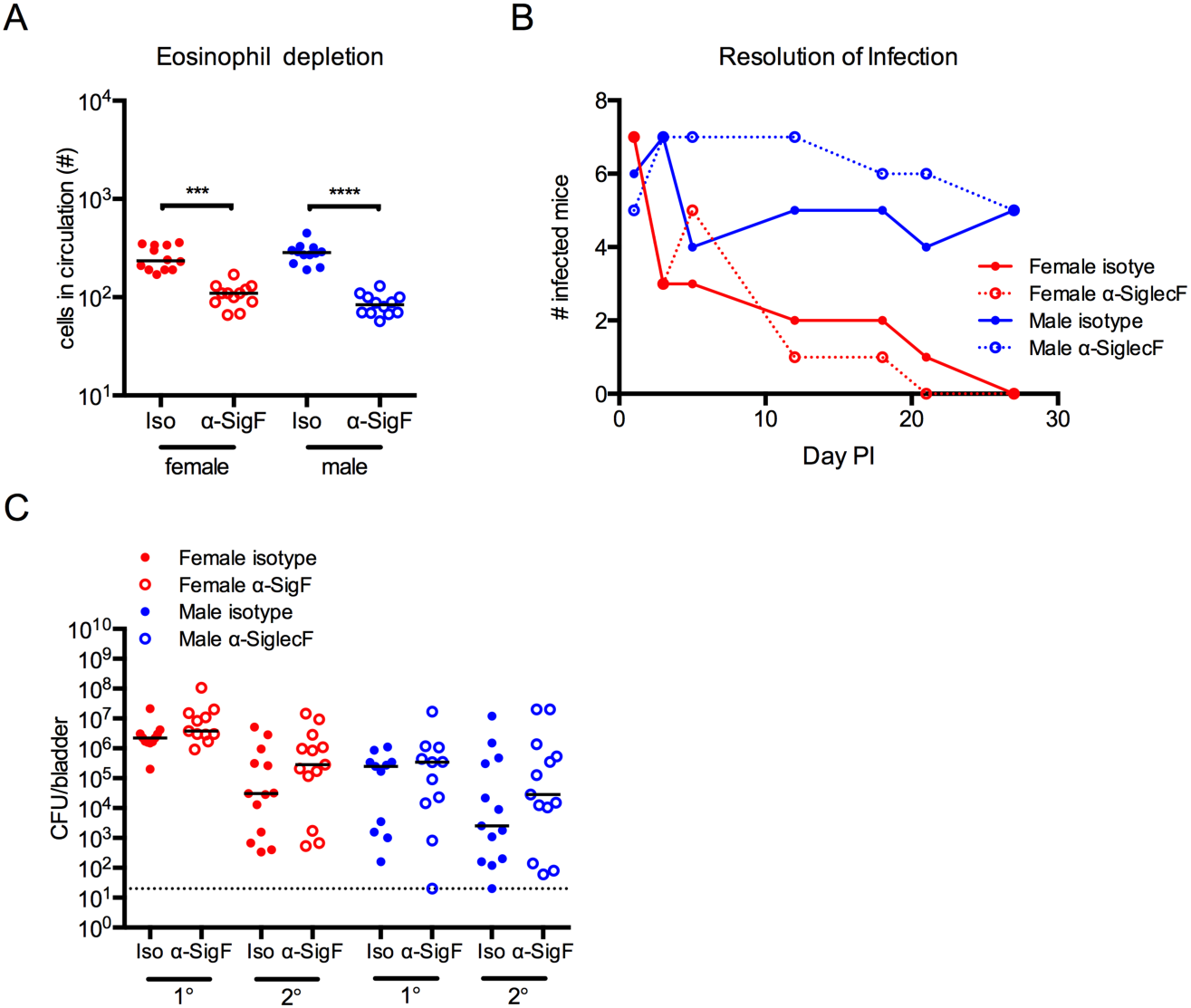
Eosinophils do not mediate resolution of UTI. (**A**-**B**) Female and male C57BL/6J mice were treated with α-SiglecF (α-SigF) antibody or isotype (Iso) control and infected with 10^7^ CFU UPEC. Graphs depict the (**A**) number of eosinophils in circulation per µL blood in control and treated mice on the day of infection, (**B**) number of infected mice determined by urine sampling over time, (**C**) CFU at 24 hours post primary (1°) or challenge (2°) infection in the bladder. In **A** and **C**, data are pooled from 2 experiments, n=6-7 mice/group in each experiment, each dot is one mouse, lines are medians. Data in **B** are representative of 3 independent experiments performed, n=6 mice/group/experiment. *** p<0.001, **** p<0.0001 Kruskal-Wallis test comparing isotype to α-SiglecF-treated mice within a single sex, with Dunn’s post-test to correct for multiple comparisons.

**Figure S5.**
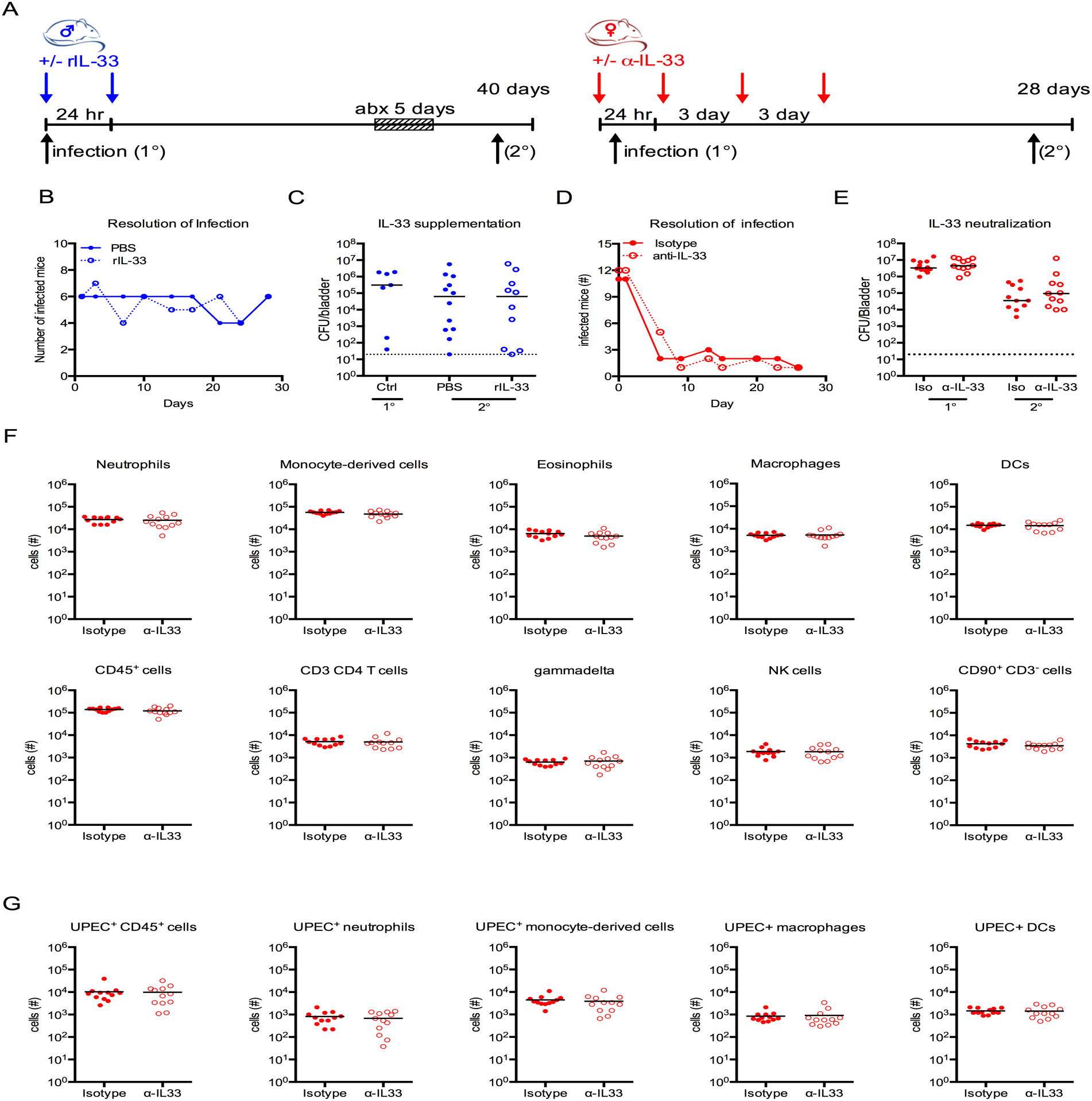
Addition or neutralization of IL-33 does not change the course of infection in male or female mice. (**A**) Experimental design in which male mice (blue) were supplemented with recombinant IL-33 (rIL-33) or female mice (red) were treated with α-IL-33 neutralizing antibody. All mice were infected with 10^7^ CFU UPEC and bacteriuria was assessed over time. (**B-G**) Graphs show (**B**) the number of infected male mice over time and (**C**) CFU at 24 hours post primary (1°) or challenge (2°) infection in the bladder, in PBS or rIL-33 treated male mice; (**D**) the number of infected mice over time and (**E**) CFU at 24 hours post primary (1°) or challenge (2°) infection in the bladder, in isotype or α-IL-33 treated female mice; and the absolute number of the (**F**) indicated immune cell populations and (**G**) UPEC^+^ immune cell populations in bladders 24 hours PI in isotype or α-IL-33 treated female mice. Data in **B** and **D** are representative experiments of 2 independent experiments, n=6 mice/group/experiment. In **C, E-G**, data are pooled from 2 experiments, n=4-5 mice/group in each experiment, each dot is one mouse, lines are medians. Mann-Whitney test, no statistically significant differences found.

**Supplemental Table 2:** Antibodies used in this study.

| **Molecule** | **Clone** | **Vendor** |
| --- | --- | --- |
| CD45 | 30-F11 | BD Bioscience |
| CD64 | X54-5/7.1 | BD Bioscience |
| CD103 | M290 | BD Bioscience |
| CD11b | M1/70 | BD Bioscience |
| CD11c | N418 | eBioscience |
| F4/80 | CI:A3-1 | AbD Serotec |
| Ly6C and Ly6G (Gr-1 antibody) | RB6-8C5 | BD Bioscience |
| Ly6G | 1A8 | BD Bioscience |
| Ly6C | AL-21 | BD Bioscience |
| MHC-II (I-A/I-E) | M5 or 114.15.2 | eBioscience |
| SiglecF | E50-2440 | BD Bioscience |
| CD3 | 145-2C11 | BD Bioscience |
| CD4 | RM4-5, GK1.5 | BD Bioscience |
| CD25 | PC61 | BD Bioscience |
| CD90.2 | 53.2.1 | ebioscience |
| IL-4Rα (CD124) | mIL4R-M1 | BD Bioscience |
| TCR $\gamma/\delta$ | GL-3 | Biolegend |
| NK1.1 | PK136 | BD Bioscience |

